# β-Ketoacyl Synthase II Homologs from a Ladderane-Producing Organism Form a Ketosynthase/Chain Length Factor-like functional heterodimer

**DOI:** 10.64898/2026.05.19.726425

**Authors:** Paola Granatino, Jolanta Nickel, Marisa I. Keller, Tobias Kutsch, Andreas Dietl, Thomas R.M. Barends

## Abstract

Ladderanes are extraordinary, highly strained chemical structures consisting of linearly concatenated cyclobutane rings. Ladderane-containing fatty acids are found in the lipids of anaerobic ammonium oxidizing (anammox) bacteria, reducing the proton permeability of their membranes to support their unique metabolism. Almost nothing is known about the biosynthesis of ladderanes, but a gene cluster unique to ladderane-producing bacteria is likely to be involved. This cluster encodes radical SAM enzymes as well as homologs of enzymes known from fatty acid biosynthesis. Amongst these are two homologs of FabF, the enzyme performing chain elongation in canonical fatty acid synthesis. The presence of two chain-elongating enzymes is unexpected; a single one would be expected to suffice, and the fact that one of the two homologs has a mutated, nonfunctional catalytic triad deepens the mystery. Here we present an in-depth characterization of the FabF homologs from anammox organisms, using biochemistry, enzymology, structural biology and computational biology. We show that the two homologs form a heterodimer analogous to the ketosynthase/chain length factor complexes known from polyketide synthases. This heterodimer is capable of performing the decarboxylation of malonate-loaded acyl carrier protein, thus initiating fatty acid biosynthesis. The crystal structure of the heterodimer explains how homodimer formation is avoided, and shows the details of the substrate binding tunnel. Mechanism-based crosslinking studies of wild-type and mutant heterodimers show the influence of residues on both subunits on substrate preference (which differs from canonical FabFs), and, together with computational studies, the crystal structure of the heterodimer in complex with substrate-loaded ACP helps explain its preference for ladderane-specific ACP. The results clearly refute an early proposal for a ladderane biosynthetic mechanism and greatly expand our current knowledge on how anammox bacteria produce their extraordinary lipids.

## Introduction

Fatty acids are essential building blocks of membrane phospholipids, in which they play multiple roles. Examples include the control of membrane fluidity through the nature of the fatty acid acyl chains (such as saturated *vs.* unsaturated, straight *vs.* branched), regulation of membrane thickness by fatty acid acyl chain length, and increase of structural integrity through the inclusion of specialized fatty acids such as mycolic acids ^1^.

Arguably one of the most interesting types of fatty acids are the ladderane fatty acids, which feature striking acyl chains with three or five cyclobutane rings linearly fused to one another, forming a ladder-like structure^2^ (Figure 1a). These fatty acids are unique to the membranes of anaerobic ammonium-oxidizing (anammox) bacteria^3^, with the most representative being the C_20_ [5]- and C_20_ [3] ladderane fatty acids, along with the homologous C_20_ [3] ladderane monoalcohols^4^ (the annotation in square brackets indicates the number of cyclobutane rings; see Figure 1a). It has been shown that phospholipids containing ladderane fatty acids reduce membrane permeability towards protons and hydroxide ions^5^. This reduction in permeability ensures the preservation of the proton gradient across the membrane of the *anammoxosome*, a specialized intracellular compartment in which the metabolism of anammox bacteria is coupled to ATP production ^6^. Given the extremely long doubling time of anammox bacteria (around two weeks), reducing proton leakage is likely essential for energy conservation and survival.

**Figure 1.**
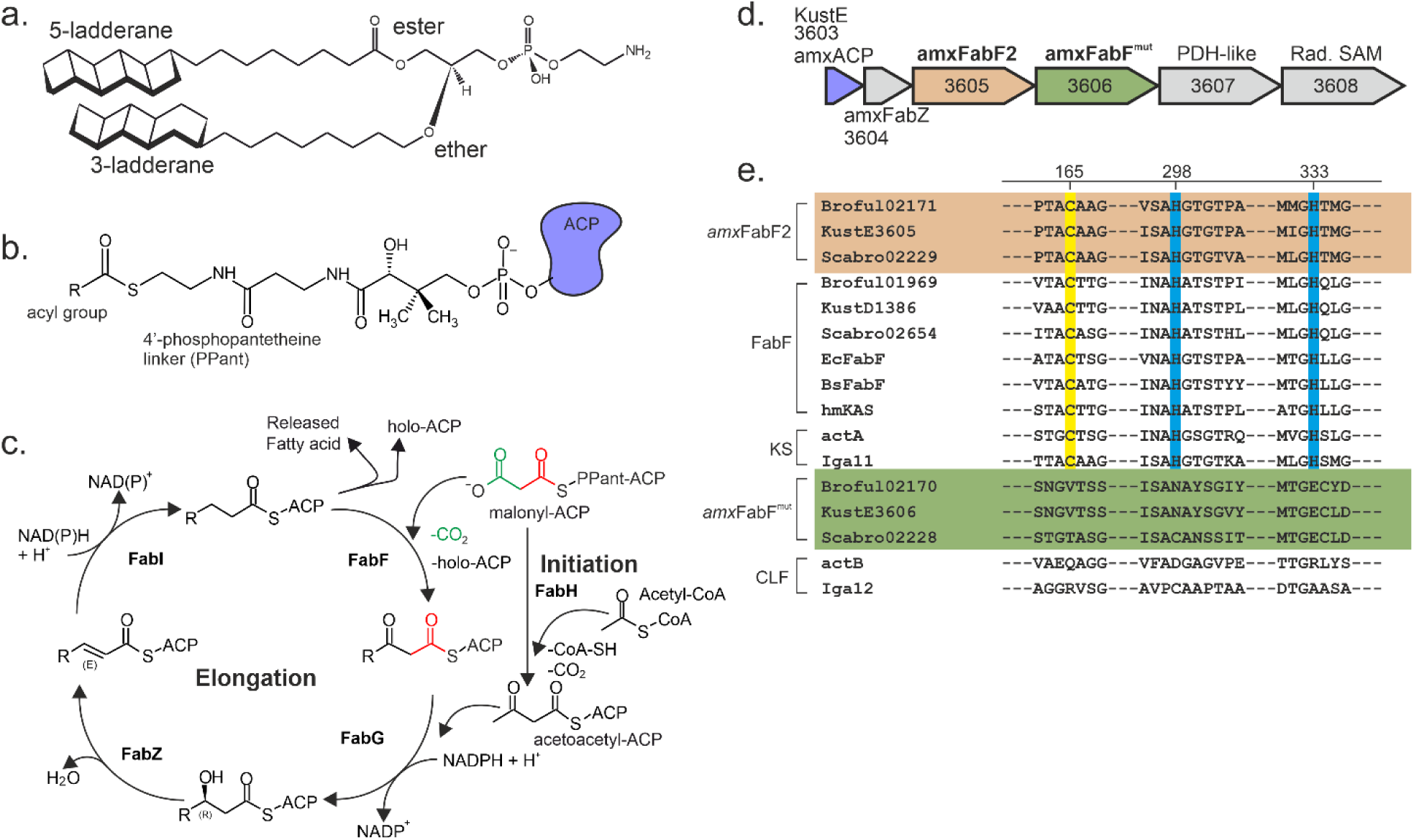
a. Example of a ladderane lipid. The glycerol phosphoethanolamine lipid depicted here contains a C_20_[5]-ladderane acyl chain connected to the glycerol backbone via an ester bond, as well as a C_20_[3]-ladderane acyl chain connected as an ether. **b.** Fatty acids are synthesized while bound to an acyl carrier protein (ACP) *via* a 4’-phosphopantetheine (PPant) linker as an acyl thioester. **c.** Canonical type II fatty acid synthesis (FAS II). A nascent fatty acid, bound to acyl carrier protein (ACP) is elongated by two carbon atoms (red) by the FabF ketosynthase using malonyl-ACP as a substrate, releasing one molecule of CO_2_. The resulting β-ketoacyl ACP is reduced by FabG, resulting in a β-hydroxyacyl ACP. This, in turn is dehydrated by FabZ and then reduced by FabI yielding an aliphatic acyl-ACP. The acyl moiety can then either be cleaved off to yield a fatty acid, or undergo another cycle of elongation. The cycle is initiated by the formation of acetoacetyl-ACP by condensing malonyl-ACP with acetyl-CoA. **d.** The putative ladderane biosynthesis gene cluster of the anammox bacterium *Kuenenia stuttgartiensis* (“Cluster I”) consists of the genes KustE3603-3608. It encodes an anammox-specific ACP (amxACP) as well as anammox-specific versions of FabZ (*amx*FabZ) and two copies of FabF (*amx*FabF2 and *amx*FabF^mut^). **e.** multiple sequence alignment, showing details of the sequences of *amx*FabF2s (beige background) and *amx*FabF^mut^s (green background) as well as of canonical FabFs, ketosynthases (KS) and chain length factors (CLF). Broful02171, KustE3605, Scabro02229: *amx*FabF2s from *B. fulgida*, *K. stuttgartiensis* and *Scalindua brodae*, respectively. Broful01969, KustD1386, Scabro02654: canonical FabFs from *B. fulgida*, *K. stuttgartiensis* and *Scalindua brodae*, respectively. EcFabF: *Escherichia coli* FabF. BsFabF: *Bacillus subtilis* FabF. hmKAS: human mitochondrial ketosynthase. actA: actinorhodin ketosynthase. Iga11: *Streptomyces sp.* ketosynthase from ishigamide biosynthesis pathway. Broful02170, KustE3606, Scabro02228: *amx*FabF^mut^s from *B. fulgida*, *K. stuttgartiensis* and *Scalindua brodae*, respectively. actB: actinorhodin biosynthesis chain length factor. Iga12: *Streptomyces sp.* chain length factor from ishigamide biosynthesis pathway. The FabF/KS active site Cys and His residues are indicated with yellow and blue boxes. The residue numbers at the top correspond to *B. fulgida amx*FabF2 numbering.

To date, it has remained unclear how ladderanes are synthesized. It has been suggested that ladderane fatty acid and -alcohols may to some extent be built analogously to canonical type II bacterial fatty acid biosynthesis (FAS II, Figure 1b, c), adapted to ladderane production by including uncommon carbon chain-modifying enzymes ^7^. Rattray and coworkers ^8^ have proposed several biosynthetic gene clusters responsible for ladderane biosynthesis, encoding both canonical FASII as well more unusual enzymes. One of these proposed gene clusters (herein called Cluster I) is both highly conserved in several anammox genera and highly expressed. This cluster (Figure 1d) encodes genes for an acyl-carrier protein (*amx*ACP), a β-hydroxyacyl-ACP dehydratase (*amx*FabZ), and two paralogues of β-ketoacyl-ACP synthase which is usually responsible for chain elongation (*amx*FabF2 and *amx*FabF^mut^). In addition to these FAS II-like proteins, Cluster I also encodes two novel enzymes, namely a putative phytoene dehydrogenase homolog (PDH) which might catalyze C-C bond desaturation, and a cobalamin-dependent radical S-adenosyl methionine (SAM) enzyme, that possibly catalyzes C-H bond functionalization for the remodeling and cyclization of the ladderane carbon skeleton. Nevertheless, there is no direct experimental evidence connecting any of the proteins encoded by Cluster I to ladderane biosynthesis, and the structures and functions of most of these proteins remain unresolved.

However, both *amx*ACP ^9^ and *amx*FabZ ^10^ have been successfully characterized and were found to exhibit some distinctive features compared to their canonical FAS II counterparts. For instance, *amx*ACP possesses an unusual structural element, a long 3_10_-helix unique to ACPs, that appears to be crucial to its functional dynamics. Moreover, *amx*FabZ displays a substrate preference very different from that of *E. coli* FabZ which preferentially acts on short and long chain acyl chains, whereas *amx*FabZ prefers acyl chains up to 6-8 carbon atoms.

Here we investigate the two paralogues of β-ketoacyl-ACP synthase encoded on Cluster I, which are analogous to FabF, the β-ketoacyl-ACP synthase II (KASII) involved in the canonical fatty acid biosynthesis pathway ^11^. Canonical FabFs are homodimeric enzymes, with each monomer contributing one active site to the dimer. FabFs build acyl chains in a stepwise fashion, adding two carbon atoms in each cycle through a ping-pong mechanism. First, a nascent acyl chain, bound to an acyl carrier protein (ACP), is transferred to a cysteine side chain on FabF in a transacylation reaction catalyzed by a catalytic triad consisting of the catalytic cysteine and two histidines (Figure 1e). The resulting acyl enzyme binds a malonyl-loaded ACP, the malonyl group undergoes a decarboxylation reaction and the remaining C_2_ moiety reacts with the acyl enzyme. This elongates the acyl chain by two carbon atoms, resulting in a β-ketoacyl product bound to ACP (Figure 1c).

If ladderane biosynthesis proceeds analogously to canonical fatty acid biosynthesis, the presence of a FabF-like enzyme on Cluster I is not unexpected, as a chain-elongating enzyme would in all likelihood be required. But why does this cluster encode two such proteins, both of which are expressed^12^And what functions might the two homologs have? In this study, we seek to address these questions using biochemical assays and biophysical- as well as structural-biological techniques.

## Materials and methods

### Expression and purification of ACP and FabF proteins

*amx*ACPs from the anammox organisms *Brocadia fulgida* and *Kuenenia stuttgartiensis* (Broful02173 and KustE3603) were expressed and purified according to previously reported protocols ^9^. For single-gene expression of *amx*FabFs, KustE3605, Broful02171 (*amx*FabF2), KustE3606, Broful02170, and Scabro02229 (*B. fulgida* and *S. brodae amx*FabF^mut^) were cloned into a pETHis-1a derived vector (a kind gift from Gunter Stier, University of Heidelberg) following amplification from genomic DNA, a kind gift from Mike Jetten, Radboud University Nijmegen. A synthetic bicistronic pRSFDuet-1-based vector containing the *amx*FabF2 and *amx*FabF^mut^ genes from *B. fulgida* was obtained from GenScript (Rijswijk, The Netherlands). The coding sequences were codon-optimized for *E. coli*, with a His-tag added to the N-terminus of *amx*FabF2. Another bicistronic vector expressing the corresponding proteins from *K. stuttgartiensis* was prepared in-house from pETHis-1a.

Recombinant *B. fulgida amx*FabF2 and *amx*FabF^mut^ proteins (from bicistronic and single-gene constructs) were expressed in *E. coli* BL21 (DE3) Codon Plus RP co-transformed with the pL1SL2 chaperone plasmid ^13^. Cultures were grown in Lysogeny Broth (LB) with 100mg/mL ampicillin, 50mg/mL kanamycin, and 30mg/mL chloramphenicol. *S. brodae* proteins were expressed in *E. coli* BL21 (DE3) without chaperone co-expression, using LB with appropriate antibiotics. Protein expression for all constructs was induced with 500μM IPTG at OD_600_∼0.6, followed by incubation at 20°C for 18 hours. Cells were harvested and resuspended in specific Lysis Buffers (see Supplementary Table 1 for all buffer compositions). Lysis was performed using a Microfluidizer (Microfluidics, USA) followed by centrifugation at 47,800×g for 1 h at 4°C in an SS-34 rotor (Sorvall SS-34 rotor, 20,000 rpm). Supernatants filtered were (0.45 μm pore size) and loaded onto Ni-NTA gravity flow columns (QIAGEN). For the purification of *B*.

*fulgida* proteins co-expressed with chaperones, columns were washed with Wash Buffer, followed by Chaperone Wash Buffer (containing ATP and KCl) to remove GroEL/ES, and washed again with Wash Buffer. Eluates were dialyzed against Chaperone Wash Buffer overnight to promote further chaperone dissociation. For all other proteins, columns were washed with high-salt Wash Buffer (200mM NaCl) and eluted immediately, and no chaperone wash steps were performed.

For the single-gene constructs, His-tags were cleaved by incubating with TEV protease during dialysis. After TEV cleavage, proteins underwent a second Ni-NTA step and were purified by size exclusion chromatography (SEC) on a HiLoad 16/60 Superdex 200 pg column (Cytiva). Final concentrations were determined using calculated extinction coefficients.

Site-directed mutagenesis was performed to introduce specific point mutations into the target expression plasmids using the QuikChange method, with PfuUltra High-Fidelity DNA Polymerase (Agilent). The oligonucleotide primers used to introduce the desired mutations are listed in Supplementary Table 3. The identity of all resulting mutant expression plasmids was verified by DNA sequencing.

### Elongation/decarboxylation assay

To directly detect the *amx*FabF2/*amx*FabF^mut^ condensation product, specifically β-keto-octanoyl-ACP, a series of *in vitro* assays were performed with subsequent analysis of the intact acyl-ACP products by LC-MS. All reactions were prepared under anaerobic conditions in a total volume of 50μL using Reaction Buffer (50 mM Tris-HCl pH 7.5, 150 mM NaCl, 10 mM MgCl_2_, 1 mM TCEP). The standard assay mixture included 3 mM ATP, 1 mM CoA-SH, 10 μM MatB, 21 μM *Ec*FabD, and 50 μM holo-*amx*ACP (KustE3603) as malonyl-ACP regeneration system^14^. The full catalytic reaction contained 10 μM of the target enzyme, *K. stuttgartiensis amx*FabF2/*amx*FabF^mut^. Four distinct conditions were tested to assess the activity of the ketosynthase components: a negative control containing no ketosynthase component, a test for the catalytic subunit *amx*FabF2 alone using 10μM *amx*FabF2, the full activity test using 10 μM of the *amx*FabF2/*amx*FabF^mut^ heterodimer, and a test using the *amx*FabF^mut^ subunit alone using 10 μM of protein from single-gene expression. Reactions were initiated by the addition of 1mM hexanoyl-*amx*ACP (loading the hexanoyl- moiety from hexanoyl-ACP by phosphopantetheine transferase, prepared as described in the Supplemental Information) and incubated at 37°C. Samples were taken at time 0 and after 1 hour. Following incubation, the reactions were quenched by the addition of 10% acetonitrile + 0.1% formic acid and analyzed directly by LC-MS for the formation of β-keto-acyl-ACP products, including acetoacetyl-ACP.

### amxACP loading with crosslinking probes

The desired bromoacyl-*amx*ACP probes (C_6_, C_8_, and C_16_) were synthesized using a one-pot enzymatic method that utilizes the substrate promiscuity of *Bacillus subtilis* phosphopantetheine transferase (Sfp). First, 2-bromo C_6_-PPant, C_8_-PPant, C_16_-PPant were prepared as described in the Supplemental Information according to protocols from ^15, 16^ and ^17^. These compounds were converted to CoA derivatives using three *E. coli* CoA biosynthetic enzymes using methods from ^18^: pantothenate kinase (CoaA), phosphopantetheine adenyltransferase (CoaD), and dephosphocoenzyme A kinase (CoaE). Subsequently, the *B. subtilis* phosphopantetheine transferase (Sfp) was employed to transfer the activated PPant-linker moiety from the CoA derivative onto the *apo*-ACP proteins, yielding the final bromoacyl-ACP probes.

The enzymatic reaction was carried out in Reaction Buffer (50 mM KH_2_PO_4_ pH 7.4, 12.5 mM MgCl2, and 1 mM DTE), adapted from Mindrebo and coworkers, 2020 ^15, 19^ using the following final concentrations: 8 mM ATP (pH 7.5), 10 μg CoaA, 10 μg CoaD, 10 μg CoaE, 40 μg Sfp, 1 mg ACP, and 200 μM PPant linker (from stocks in DMSO; 100 mM for the 2-bromo C_6_- and C_8_-PPant, 10 mM for 2-bromo C_16_-PPant).

Reactions were incubated at 37°C overnight, followed by purification via size exclusion chromatography (SEC) using a HiLoad Superdex 75 (16/60) column. The SEC running buffer (GFB) was 150 mM NaCl, 50mM HEPES, pH 7.5. Eluted protein fractions were collected and concentrated using Amicon Ultra Centrifuge tubes with a 3 kDa molecular weight cut-off (MWCO). Probe concentration was calculated based on UV absorbance measurements at 280 nm using a theoretical extinction coefficient *amx*ACP (ignoring any contribution from pPant).

### Crosslinking assay

The substrate specificity of the wild-type *amx*FabF2/*amx*FabF^mut^ heterodimer and its mutants was assessed using a mechanism-based crosslinking assay adapted from Mindrebo et al. ^15, 19^. Reactions were set up in Assay Buffer (25 mM Tris pH 7.4, 150 mM NaCl, 10% (v/v) glycerol, and 0.5 mM TCEP). Each reaction contained the purified ketosynthases (wild-type or mutant) at a final concentration of 2 μM and the respective 2-bromoacyl-*amx*ACP probe at a final concentration of 4 μM. The reactions were incubated at room temperature. At designated time points (0, 1, 5, 15, 30, 60, 180, 480 min), a 10 μL aliquot was removed and quenched by mixing with 6 μL of 4×SDS loading dye. The resulting samples were immediately analyzed by 12% SDS-PAGE. Gels were stained with InstantBlue® Coomassie Protein Stain (Abcam), and the intensity of the protein bands corresponding to the unreacted *amx*FabF2 and the crosslinked adduct was quantified using densitometry with ImageJ^20^. The crosslinking efficiency at each time point was calculated as the percentage of the crosslinked adduct relative to the total amount of *amx*FabF2. All experiments were performed in biological triplicates. The final plotted data represent the average normalized crosslinking efficiency, with error bars indicating the standard deviation (±SD).

### Crystallization, diffraction data collection, structure solution, and refinement

*S. brodae amx*FabF^mut^, B*. fulgida amx*FabF2/*amx*FabF^mut^ and the *B. fulgida amx*FabF2/*amx*FabF^mut^ -*amx*C_8_ACP complex were concentrated to ∼10 mg/ml in 25 mM HEPES/KOH pH 7.5, 100 mM KCl using Amicon Concentrators. In the case of the *amx*FabF2/*amx*FabF^mut^ -*amx*C_8_ACP complex, 0.5 M arginine, pH 6.5 was added to the buffer. Initial crystallization screens were performed using a Mosquito pipetting robot (SPT LabTech Ltd., Melbourne, UK), using 100+100 nl drops in Greiner XTL Low Profile 96-well crystallization plates (Greiner Bio One, Frickenhausen, Germany) equilibrating against 70 μl of reservoir solution from commercial crystallization screens (JSCG Core I-IV, PEG Suite, Classics Suite, Ammonium Sulfate Suite, ProComplex Suite, Quiagen, Hilden, Germany, as well as Wizard I and II (Emerald, Bainbridge Island, USA), complemented as needed with Additive Screens 1-3 (Hampton Research, Alison Viejo, USA). Initial hits were optimized in 24-well crystallization plates (ICN Biomedicals Inc., Aurora, USA) using 1+1 μl hanging drops equilibrating against 700 μl reservoir solution. Diffraction data were collected remotely by Ilme Schlichting and M. Tarnawski at the PX-II beamline of the Swiss Light Source (Villigen, Switzerland) or at the ID23-EH1 beamline of the ESRF (Grenoble, France) and processed using the automatic processing pipelines at these facilities using XDS^21^ and POINTLESS^22^. Data collection and refinement statistics are listed in Supplementary Table 2.

Data sets were phased by molecular replacement using PHASER^23^ as implemented in the PHENIX package^24^. Search models were taken from the PDB or generated using the AlphaFold2^25^ implementation provided by ColabFold^26^. Map inspection and rebuilding were done using COOT^27^ and refinement was performed using PHENIX. Structural figures were prepared using PyMOL^28^. The stereochemical quality of the model was validated using MolProbity^29^.

### Electrostatic surface potential calculation

Electrostatic surface potentials were calculated using the Adaptive Poisson-Boltzmann Solver (APBS) ^30^ plugin within the PyMOL Molecular Graphics System. Prior to calculation, protein structures were prepared by removing all non-protein atoms and assigning protonation states appropriate for a pH of 7.0 using the PDB2PQR software server. The resulting electrostatic potential was mapped onto the molecular surface and visualized, with potentials displayed on a color scale from -5 kT/e (red) to +5 kT/e (blue).

## Results and discussion

### Sequence comparison and prediction of function

We aligned the sequences of the two FabF homologs encoded by Cluster I, *amx*FabF2 and *amx*FabF*^mut^* , with sequences of canonical homodimeric β-keto-acylACP ketosynthases (*E. coli* and *Bacillus subtilis* FabF) from type II fatty acid synthase (FASII; White 2005) as well as of canonical FabFs from different anammox bacteria (Figure 1e, Supplemental Figure S1). The alignment shows that the two FabF paralogues share 27-32% sequence similarity with canonical FabFs. The active sites display particularly high conservation of the cysteine/histidine/histidine (Cys-His-His) catalytic triad (White, 2005), which are different only in amxFabF^mut^. According to the generally accepted catalytic mechanism of FabFs, the conserved cysteine at the *amx*FabF2 monomer position 165 (using *Brocadia fulgida* numbering) would accept the growing fatty acid chain, whereas the histidines at position 298 and 333 would make up the rest of the catalytic triad. However, as stated before, this Cys-His-His triad is absent in the *amx*FabF^mut^ proteins.

Rattray *et al.*^8^ predicted that these mutations would lead to correctly folded, but inactive enzyme, *i.e.* a protein that is unable to catalyze either decarboxylation of malonyl-ACP or chain elongation. In *amx*FabF^mut^ from most anammox species, the conserved catalytic cysteine is replaced by a valine, except in *Scalindua brodae* where it is replaced by a threonine (Figure 1e, Supplemental Figure S1). However, either mutation would abrogate the normal acyl chain elongation function of a ketosynthase. Additionally, the two histidines of the catalytic triad are mutated to asparagine, glutamate or cysteine in *amx*FabF^mut^s (Figure 1e, Supplemental Figure S1), all of which would severely hinder catalytic activity even if the catalytic cysteine were retained.

Interestingly however, the organization of the genes coding for *amx*FabF2 and *amx*FabF^mut^, *i.e.* with *amx*FabF^mut^ encoded just upstream from *amx*FabF2, resembles the organization of the highly homologous ketosynthase (KS) and chain-length factor (CLF) genes in some type II polyketide biosynthesis systems (PKSs), which are also often localized in the same cluster^31^.

These ketosynthases are enzymes that are evolutionarily highly related to FabFs and catalyze essentially the same reaction, elongating a polyketide chain bound to a carrier protein, using carrier-protein-bound malonate as a building block. However, they do not form homodimers, but heterodimers of an active ketosynthase bound to an inactive homolog called a “Chain Length Factor” (CLF). This CLF controls ketosynthase product chain length through its influence on the substrate binding tunnel formed at the interface of the heterodimer. Importantly, like *amx*FabF^mut^, these CLFs do not contain a typical Cys-His-His catalytic triad, as *e.g.* in the CLFs Iga12 ^32^ and ActB^33^ from the ishigamide and actinorhodin PKS systems, respectively, which are also depicted in the sequence alignment in Figure 1e and Supplemental Figure S1. Thus, we hypothesized that *amx*FabF2 and *amx*FabF^mut^ form a functional heterodimer reminiscent of a PKS ketosynthase / chain length factor heterodimer. In this proposed heterodimeric complex, *amx*FabF2, with its intact catalytic triad, would function as the decarboxylating/condensing ketosynthase enzyme, while *amx*FabF^mut^ could act as a chain length factor, regulating the chain length of the products. We therefore set out to test this hypothesis.

### *amx*FabF2 and *amx*FabFmut display decarboxylation activity together but not alone

The anammox model organism *Kuenenia stuttgartiensis* Cluster I *amx*FabF2 (gene locus *KustE3605*) and *amx*FabF^mut^ (*KustE3606*) were heterologously expressed in *E. coli* and purified as separate constructs. Mass spectrometry (MS)-based *in vitro* assays were carried out to evaluate if the proteins were enzymatically active. The proteins were incubated with their potential substrates, malonyl-*amx*ACP as starting unit and hexanoyl-*amx*ACP (*KustE3603*) as the substrate to be elongated, to see whether the proteins displayed any condensation activity by producing β-ketooctanoyl-ACP or decarboxylation activity on malonyl-*amx*ACP. The reaction mixtures were analyzed *via* High Performance Liquid Chromatography (HPLC) coupled to electrospray ionization (ESI) MS. Neither protein displayed detectable activity of any kind under the tested reaction conditions.

The two proteins were then co-expressed, with *amx*FabF2 tagged with an N-terminal His-tag for affinity purification, while *amx*FabF^mut^ displayed no tag. During the purification steps using Ni-NTA (nickel- nitriloacetic acid) affinity chromatography, the two proteins co-eluted, indicating heterodimer formation (Supplemental Figure S2). Functional assays of the co-expressed proteins on a mixture of malonyl- and hexanoyl-ACP revealed decarboxylation activity towards malonyl-ACP yielding acetyl-ACP, but no chain elongation could be detected. Interestingly, a peak corresponding to the mass-to-charge ratio (*m*/*z*) of the β-keto derivative acetoacetyl-ACP, *i.e.* the compound with which fatty acid synthesis is initiated (Figure 1), was visible in the spectra under the tested reaction conditions. This suggests that the heterodimer is able to catalyze the initial condensation step in the FAS II process, formation of acetoacetyl-ACP, even if further elongation cycles were not observed under these *in vitro* conditions. We named this heterodimeric complex of the two homologs *amx*FabF^het^.

#### *amx*FabF^mut^ crystal structure

To obtain structural information on Cluster I FabF homologs, we first attempted to crystallize both *amx*FabF2 and *amx*FabF^mut^ from various anammox organisms separately, expressed as single gene constructs. *amx*FabF2 did not yield crystals for any construct tried. However, we were able to obtain crystals of *amx*FabF^mut^ from the anammox organism *Scalindua brodae* which diffracted to 2.2 Å resolution. A data set was collected and phased by molecular replacement using the *B. subtilis* FabF structure (PDB: 4LS6) as the search model. The asymmetric unit of the final, refined structure (see Supplementary Table 2 for data- and model statistics) contains two homodimers of a similar homodimeric structure as other FabFs (Figure 3a.). The thiolase-fold structure shared by the four monomers is very similar to that of *B. subtilis* FabF, indeed, that protein is identified as the closest structural homolog of *S. brodae amx*FabF^mut^ in the PDB by the DALI^34^ server. Residues 110 to 130 did not show sufficient electron density for model building in any of the monomers in the asymmetric unit and this region is probably disordered. In canonical FabFs, these residues are part of a helix-loop-helix motif formed by residues 110-140 that is involved in dimerization by interacting with a helix on the other monomer. Importantly, in FabFs the helix-loop-helix motifs from the two monomers normally cover the substrate binding sites.

**Figure 2.**
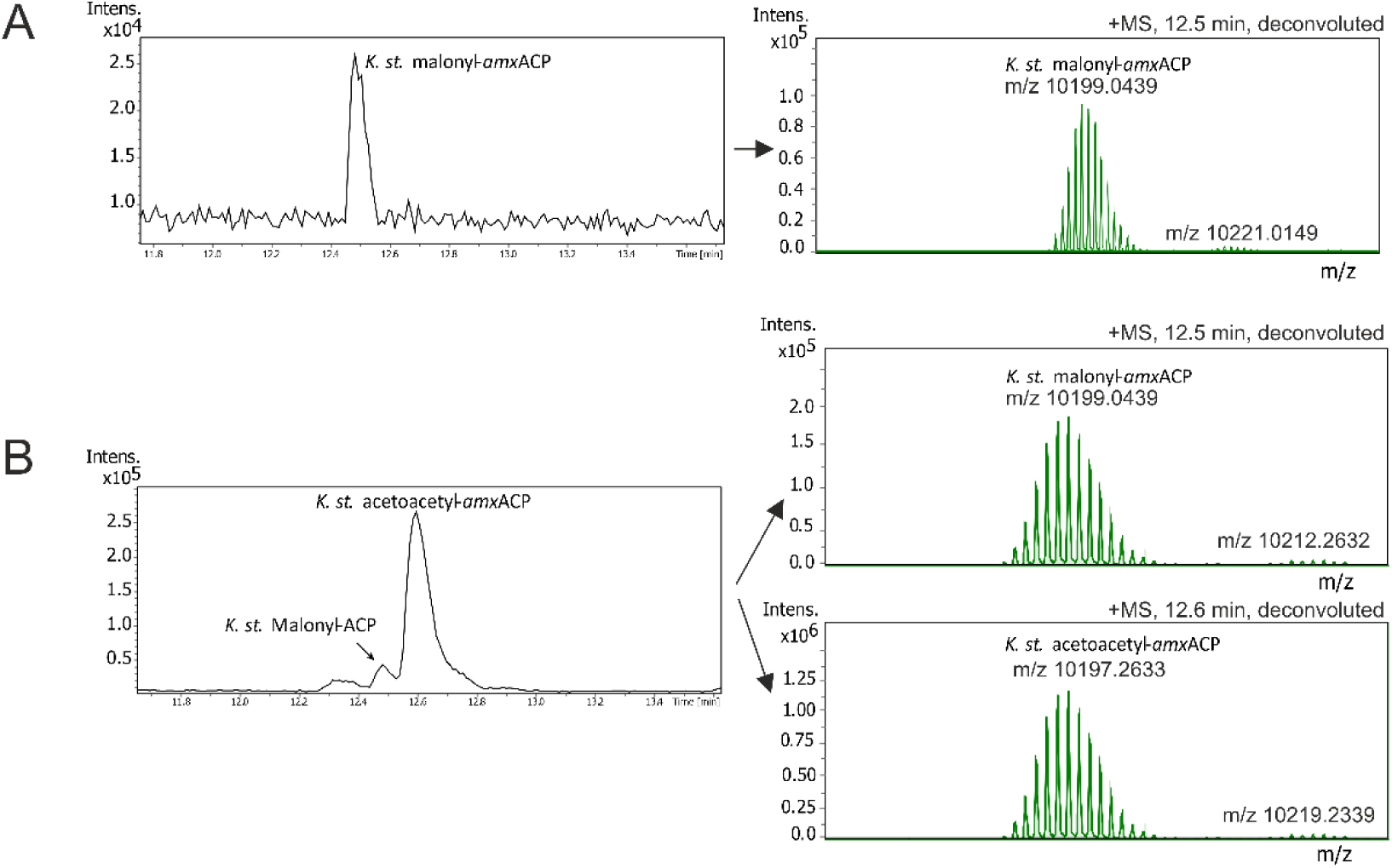
A. Left panel: LC-MS trace of a mixture containing malonyl- and hexanoyl-loaded *amx*ACP but no FabF homologs zoomed in on the region where malonyl-*amx*ACP elutes. Right panel: deconvoluted mass spectrum of the malonyl-*amx*ACP. **B.** LC-MS trace of the same mixture incubated with co-expressed *amx*FabF2 and *amx*FabF^mut^. The malonyl-*amx*ACP peaks has almost disappeared and a peak for acetoacetyl-*amx*ACP has appeared. right: deconvoluted mass spectra of the peaks in the left panel.

**Figure 3.**
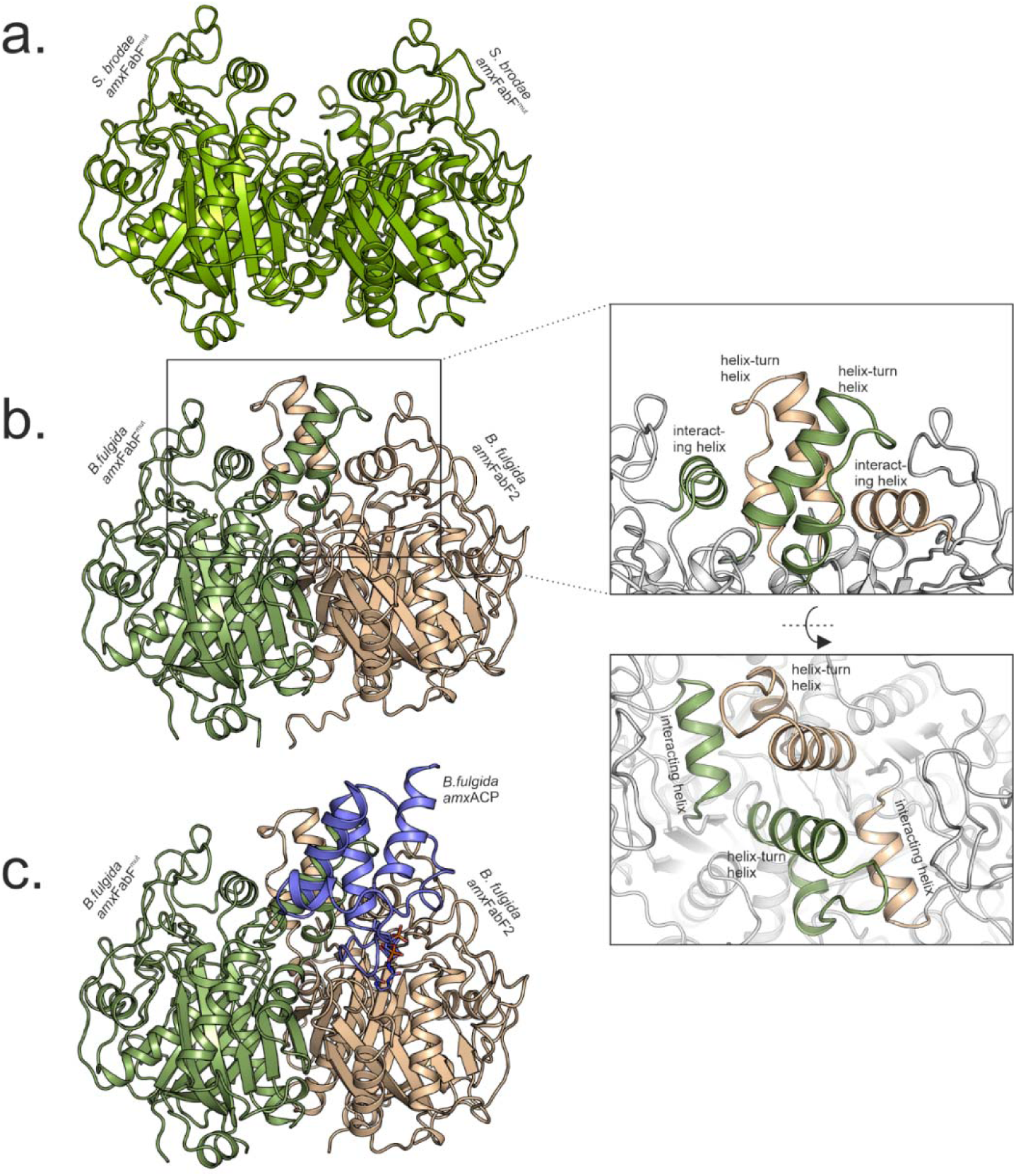
Crystal structures of anammox FabF homologs. **a.** Homodimer of *Scalindua brodae amx*FabF^mut^. **b.** *amx*FabF^het^ heterodimer consisting of monomers of *Brocadia fulgida amx*FabF2 (beige) and amxFabFmut (green). The insets focus on the helix-turn-helix motifs and the interacting helices from both monomers; the rest of the protein is shown in green. The lower inset shows the structure rotated by 90 degrees. Note that no electron density is visible for the helix-turn-helix motifs in the *amx*FabF^mut^ homodimer shown in panel a. **c.** *B. fulgida amx*FabF^het^ functionally crosslinked with *B. fulgida amx*ACP.

#### Crystal structure of the *amx*FabF heterodimer

The *Brocadia fulgida amx*FabF^het^ structure was determined at 2.6 Å resolution (see Supplementary Table 2 for data- and model statistics). The heterodimeric structure (Figure 3b) is composed of one *amx*FabF2- and one *amx*FabF^mut^ subunit, with both monomers showing the typical fold of a FabF monomer. Interestingly, the closest structural homolog of the *amx*FabF2 subunit as found by DALI ^34^ is not a FabF but the ketosynthase from the ishigamide biosynthesis gene cluster (pdb 6kxe, ^32^) with an RMSD of 1.1 Å for 395 aligned Cαs. For the *amx*FabF^mut^ subunit, however, the closest structural homolog is the FabF from *B. subtilis* (pdb 4ls5, ^35^) with an RMSD of 1.7 Å for 392 aligned Cαs. The overall structure of each monomer consists of an α-β-α-β-α thiolase fold core, with a series of α-helices and loops on one side. In the active site in *amx*FabF2, the catalytic triad assumes the typical conformation known from other FabFs. A magnesium ion is bound close to the three catalytic residues, coordinated by the backbone atoms of Ser296, Ala297, and Asn390, as well as the side chains of Ser296, Glu344, and Asn389, all from *amx*FabF2. Huang *et al.* ^36^ showed that sodium or potassium ions bind at a similar site in *Helicobacter pylori* FabF, and that the presence of a metal ion at this position is critical for the enzymatic activity in that system, primarily by improving the structural stability of the enzyme.

Importantly, the region consisting of residues 110 to 140, which as mentioned forms a motif involved in dimerization and substrate binding site formation in FabFs, and which was largely disordered in the *amx*FabF^mut^ structure, is clearly defined for all *amx*FabF2 and *amx*FabF^mut^ molecules in the maps of the heterodimeric *amx*FabF^het^ structure. Dimerization results in the formation of two long tunnels in the heterodimer, but these tunnels differ strongly from each other in *amx*FabF^het^, rather than being identical as in the homodimeric, canonical FabFs. One runs from an entrance on the surface of *amx*FabF2 *via* the active site with the catalytic Cys-His-His triad towards the dimer interface splitting into a long- and a short branch, whereas the other one runs from an entrance on the *amx*FabF^mut^ monomer surface towards the inactive, mutated triad in *amx*FabF^mut^ and from there further into the heterodimer (Figure 4). In the crystal structure, this latter tunnel harbors a PEG molecule from the crystallization solution and is unlikely to be involved in catalysis, having no access to the only catalytic triad on *amx*FabF2. The other tunnel, starting on the surface of *amx*FabF2 does pass through the active site, and is therefore likely the substrate binding tunnel. Closest to the entrance, the wall of this tunnel is hydrophilic, which is expected, as this is where the phosphopantetheine linker between *amx*ACP and the substrate (Figure 1) would need to bind; in the crystal structure, a HEPES molecule from the crystallization solution binds here. The rest, however, is mainly surrounded by hydrophobic residues: Ala110, Ile112, Ala 164, Phe203, and Met337 from the *amx*FabF2 monomer, as well as Phe135, Thr138, and Val139 from the *amx*FabF^mut^ subunit cover the long and short branches. To visualize what substrates could fit inside this tunnel system, we modeled linear, saturated fatty acids with their C1 carbon atoms in close proximity to the active Cys165 as would be expected for an enzyme-substrate complex. This shows that a C_16_ fatty acid would fit into the long branch, whereas the short branch would fit a C_12_ fatty acid (Figure 4).

**Figure 4.**
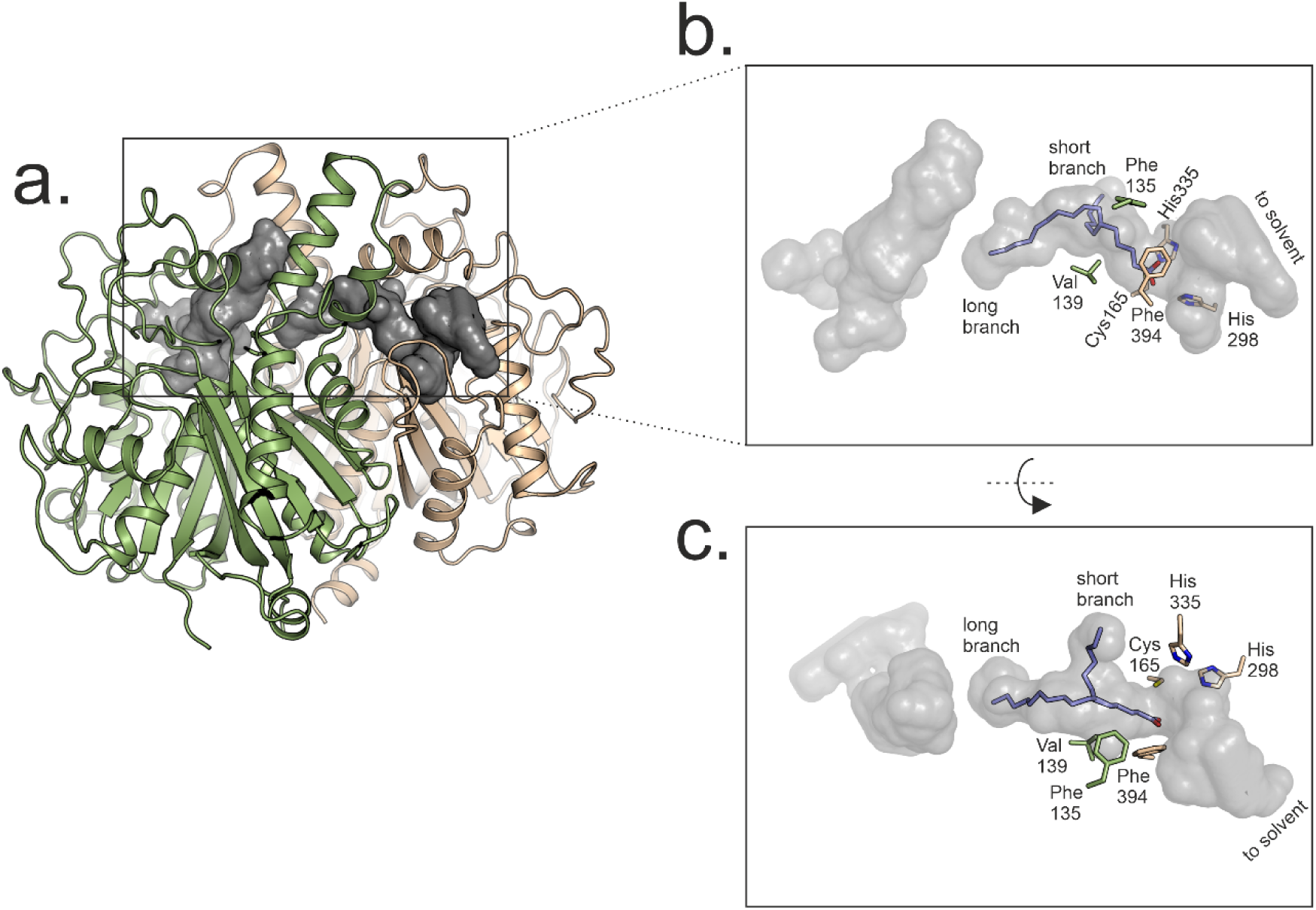
Tunnels in *amx*FabF^het^. **a.** Overview of the dimer, with *amx*FabF2 in beige and *amx*FabBmut in green. The tunnels are shown in grey. **b.** and **c.** Closeup of the tunnel systems in two perpendicular orientations, with C_16_ and C_12_ fatty acids modeled into the long and short branch of the tunnel visiting the active site. The active site Cys165, His298 and His335 are shown as sticks, as are Phe394 from *amx*FabF2 and Phe135 and Val139 from the helix-loop-helix motif from *amx*FabF^mut^.

## Electrostatic Surface Analysis Reveals an Asymmetric Dimer Interface in *amx*FabF^het^

To investigate the interactions between *amx*FabF2 and *amx*FabF^mut^ in the heterodimer, we calculated their electrostatic surface potentials using APBS ^30^ and compared them to that of the monomers of the canonical *E. coli* FabF homodimer. This revealed major differences, particularly concerning the helix-loop-helix motifs (residues 110-140). As mentioned before, the helix-loop-helix motif is present in both subunits and interacts with a helix on the respective other subunit that consists of residues 196-206 in both subunits. In the *E. coli* FabF homodimer, the interfaces between the helix-loop-helix motifs and their interacting helices are necessarily symmetrical and comprise predominantly hydrophobic interactions. In stark contrast, the corresponding interfaces in amxFabF^het^ reveal a pronounced electrostatic asymmetry; the helix-loop-helix motif and interacting helix of *amx*FabF2 display a clear negative surface potential, whereas the helix-loop-helix motif and interacting helix on *amx*FabF^mut^ have a distinctly positive surface potential (Supplemental Figure S3). This electrostatic asymmetry provides a clear structural rationale for the heterodimerization of *amx*FabF2 or *amx*FabF^mut^ into *amx*FabF^het^, as hypothetical homodimers of either subunit would be destabilized by charge repulsion. This likely also explains why the helix-loop-helix motif was mostly unresolved in the crystal structure of the *amx*FabF^mut^ homodimer from *Scalindua brodae*, as the repulsion between the helix-loop-helix motifs and their interacting helices would preclude these from forming specific interactions, leading to their being disordered.

### Substrate preference of the *amx*FabF^het^ heterodimer

Our initial MS based *in vitro* assays did not confirm whether the *amx*FabF^het^ heterodimer could process its potential substrates to yield longer chain products than acetoacetyl-ACP. We therefore looked for other methods to probe the substrate binding profile of the *amx*FabF^het^ heterodimer. In particular, given the possibility that *amx*FabF^mut^ acts as a chain-length factor, we wanted to investigate the specificity of the heterodimer towards substrates with various acyl chain lengths. To that end, we employed mechanism-based crosslinking probes developed by Burkart and colleagues^16, 37, 38^. Specifically, we prepared α-bromoacids of various lengths (C_6_, C_8_, and C_16_, supplemental Figure S4) and loaded them onto *amx*ACP. When an ACP loaded with such a probe reacts with a catalytically competent FabF active site, the active site cysteine will attack the α-bromo acyl group, resulting in the formation of a covalent FabF=acylACP adduct that can be quantified using SDS-PAGE (supplemental Figure S5. In this way, the propensity of the FabF being studied to interact with ACPs loaded with probes containing acyl chains of different natures can be investigated^16, 37, 38^.

The synthesis of these probes, as detailed in the Supplementary Information, first involved the production of a pantothenic acid analogue. This was then linked to α-bromo acids of various lengths *via* a non-hydrolysable amide bond. The resulting compounds were converted to phospho-coenzyme A (CoA) derivatives using the *E. coli* CoA biosynthetic enzymes pantothenate kinase, phosphopantetheine adenyltransferase, and dephosphocoenzyme A kinase. The thus activated, acylated phosphopantetheines were then loaded onto *amx*ACP by the *B. subtilis* phosphopantetheinyl transferase Sfp (Supplemental Figure S6). Correct *amx*ACP loading was confirmed by electrospray-ionization MS.

We first investigated the reactivity of the individual components of the heterodimer towards *amx*ACP loaded with these probes using a single-time point crosslinking assay. *B. fulgida amx*FabF2 and *amx*FabF^mut^ from independent single gene expression were incubated with C_6_, C_8_, and C_16_ *amx*ACP (also from *B. fulgida*). After a 24-hour incubation period, no reaction of any of the ACP probes with either of the two separate protein subunits could be detected. Then, heterodimeric *amx*FabF^het^ was incubated with an excess of probe-loaded *amx*ACP and the formation of any *amx*FabF=acyl-*amx*ACP adduct was monitored over a 24-hour time course, with samples collected at multiple time points and analyzed *via* SDS-PAGE (Supplementary Figures S7-S9). Densitometric analysis was used to quantify the relative amounts of the *amx*FabF=acyl-*amx*ACP complex, visible on the gel as the *amx*FabF2=*amx*ACP adduct. The relative intensities were plotted against the respective time points for each sample (Figure 5).

**Figure 5.**
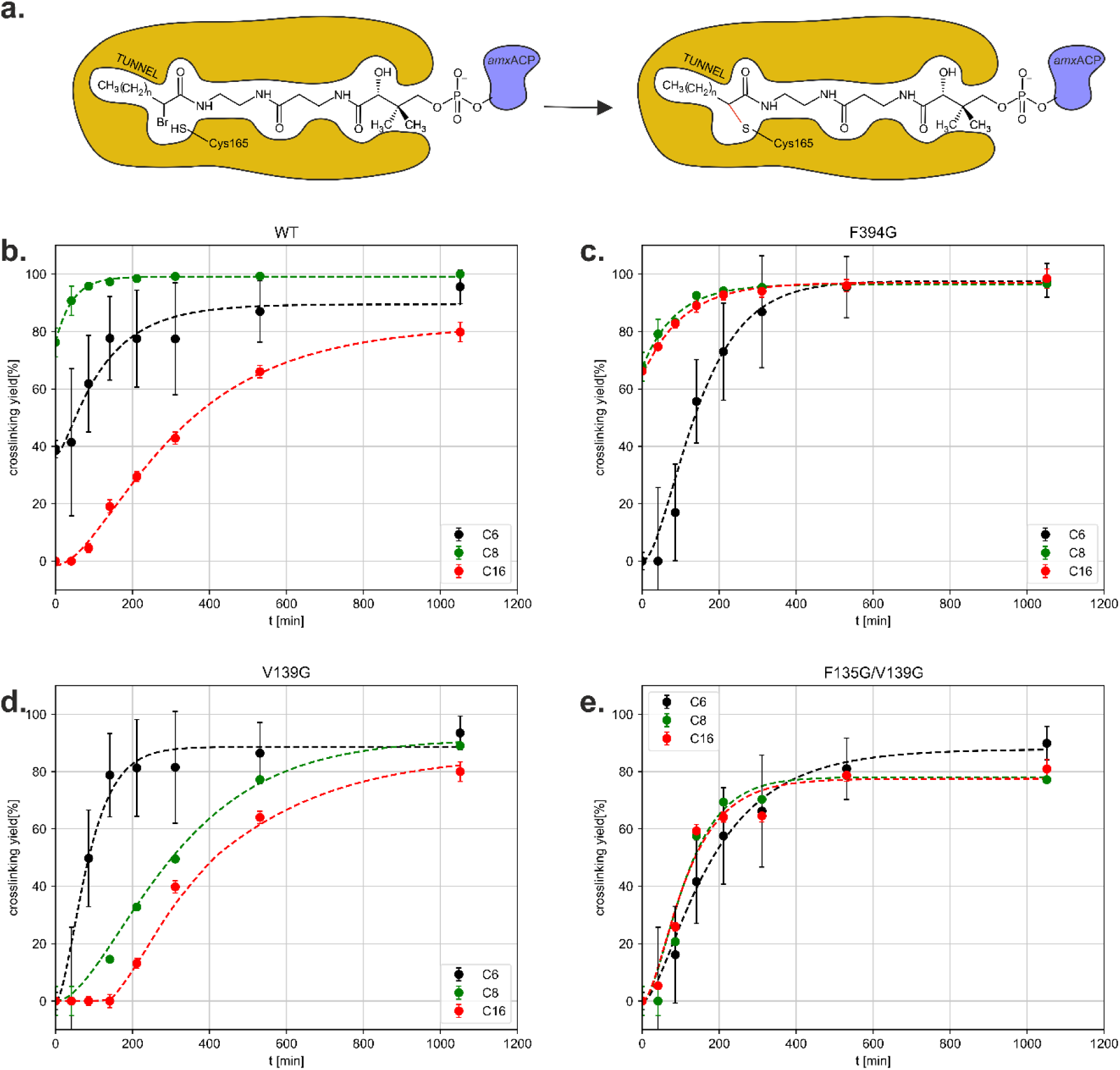
a. Principle of the crosslinking assay. **b.-e.** Fraction of *amx*FabF2 WT and mutants crosslinked as a function of time in reactions with C_6_- (black lines), C_8_- (green lines) and C_16_-probe-labeled *amx*ACP (red lines)

In samples containing *amx*FabF^het^ and a twofold molar excess of C_8_-acyl-ACP probe, complete depletion of free *amx*FabF2 and formation of the *amx*FabF2=acyl-*amx*ACP complex was observed in about three hours. Using the C_6_ probe, the experiment showed poor reproducibility, but it was clear that the crosslinking reaction proceeded more slowly and did not reach completeness. With the long-chain C_16_ substrate, however, formation of the *amx*FabF2=acyl-*amx*ACP complex was only detectable after a 60-minute lag phase, and the reaction also did not reach completion even after 24 hours.

We then mutated residues in positions known to influence substrate binding in ketosynthase/chain length factor complexes and repeated the crosslinking assays. Specifically, we mutated Phe394 on *amx*FabF2 and Val139 and Phe135 on the helix-loop-helix motif of *amx*FabF^mut^, all of which line the substrate binding tunnel (Figure 4). Phe394 is known as a “gate” residue, whose sidechain can assume conformations that either block or allow access to the part of the binding tunnel where the substrate’s acyl chain should bind. When this residue on *amx*FabF2 was mutated to glycine, crosslinking assays with all probes reached completion, with the reaction with the C_8_ probe proceeding slightly more slowly than in the case of WT protein. Moreover, the reaction with the C_16_ probe proceeded essentially as quickly as the reaction with the C_8_ probe. Reactions with the C_6_ probe, however, started more slowly than with the WT protein. We then turned to Val139 on *amx*FabF^mut^, which lines the wall of the substrate binding tunnel approximately where the C8 atom of a substrate would bind (Fig. 4b,c). The corresponding residue (a leucine) in human mitochondrial ketosynthase was found to restrict binding of substrates longer than C_8_ ^39^. When Val139 was mutated to glycine in our heterodimer, the reaction proceeded most quickly with the C_6_ probe, whereas the reaction with the C_8_- over the C_16_ probes showed a pronounced lag phase. Next, we investigated Phe135, also on *amx*FabF^mut^, which constricts the width of the acyl chain binding part of the tunnel, also close to where a substrate’s C8 atom would bind. The Phe135Gly single mutant did not express, but with the Phe135Gly/Val139Gly double mutant, the reactions with all probes showed essentially the same kinetics. With neither the Val139Gly nor the Phe135Gly/Val139Gly mutant, however, did any of the reactions reach completion in 24 hours. Taken together, while the crosslinking reaction used here is not the physiological reaction catalyzed by *amx*FabF^het^, the results do suggest that residues from the inactive *amx*FabF^mut^ affect the substrate length preference of the heterodimeric complex as expected for a ketosynthase/chain length factor complex.

### Crystal structure of *amx*FabF^het^ in complex with C8-ACP

To further investigate the interaction between *amx*FabF^het^ and substrate-loaded *amxACP*, we prepared the *Brocadia fulgida amx*FabF^het^-C_8_-*amx*ACP complex using a C_8_ 2-bromo pantetheinamide probe as a cross-linker and crystallized it. The 2.1 Å resolution crystal structure of this ternary complex was determined by molecular replacement starting from the crystal structure of the heterodimer (see Supplementary Table 2 for data- and model statistics). The asymmetric unit contains two *amx*FabF^het^ dimers, each of them showing clear density for the PPant-acyl linker between *amx*FabF2 and *amx*ACP, but for only one the density of the *amxACP* molecule was sufficiently well defined to build it, suggesting that in the other complex it is disordered.

In the fully resolved complex, *amx*ACP contacts both *amx*FabF*2* and *amx*FabF^mut^ subunits, with interaction surfaces of approximately 250 Å² and 300 Å², respectively. On *amx*ACP, the interactions occur predominantly *via amx*ACP helix II (from Ser41 to Ala49), which contacts the loop part of the helix-loop-helix motif on *amx*FabF2, mirroring the interactions noted in the in the *E. coli* FabZ-ACP and FabF-ACP complexes ^40, 41^ as well as in the *Helicobacter pylori* ACP-FabF complex^36^. However, in the *amx*ACP-*amx*FabF^het^ structure, the orientation of the acyl carrier protein’s four-helical bundle has the helices oriented more perpendicularly towards the enzyme’s surface than in the *Helicobacter* and *E. coli* structures. This was also observed in the *amx*ACP-*amx*FabZ dehydratase complex^10^ and is likely a property of *amx*ACP.

To further investigate the recognition between *amx*ACP and *amx*FabF^het^, we performed Monte-Carlo interaction studies using the program MCMAP ^42^, which searches for favorable interactions between protein molecules based on electrostatics, and allows the prediction of interaction surfaces for even very fleeting interactions such as the encounter complexes between electron donor and -acceptor proteins ^42^ ^43^. As the MCMAP method is based on electrostatics, we hoped it might be applicable to interactions of acyl carrier proteins with fatty acid biosynthesis enzymes, which often involve electrostatic complementarity. Indeed, when used to search for interaction modes between *amx*ACP and *amx*FabZ it correctly predicted all the interaction surfaces between the two proteins (see Supplemental Figure 4). The same was true when used to predict interactions between *H. pylori* ACP and -FabF. We therefore used it to predict interactions between *amx*FabF^het^ and *amx*ACP. This found three preferred locations for *amx*ACP around *amx*FabF^het^ (Supplemental Figure S10). Of these, the most populated one corresponds to the position of the carrier protein in the crystal structure of the complex, i.e., close to the “positive patch” on the surface of *amx*FabF^het^ at the entrance to the active site in *amx*FabF2. The other two, one of which is minor, are close to other positively charges sites on the surface of *amx*FabF^het^.

Interestingly, MCMAP does not predict preferential location of *amx*ACP close to the entrance of the “defunct” tunnel in *amx*FabF^mut^. This is in contrast to the situation in the homodimeric *Helicobacter* FabF. There, the program also predicts three preferred positions for ACP around the FabF homodimer but two of these are at the entrances to the tunnels of the two active sites, and the third one is at a distant positively charged surface patch. These results suggest that the surface properties of the two FabF homologs in *amx*FabF^het^ have evolved to favor substrate binding at the entrance to the only active site in the heterodimer over binding at the tunnel leading to the mutated active site in *amx*FabF^mut^. Moreover, the program predicts that, on the side of *amx*ACP, a group of negatively charged residues at one side of *amx*ACPs four-helical bundle is involved in recognition, rather than helix II as is usually the case for ACPs (Supplemental Figure XXX). This might contribute to specificity for *amx*ACP over any of the other ACPs present in *Brocadia*.

Indeed, *Scalindua brodae amx*FabZ was found to be specific for substrates bound to *amx*ACP over those bound to the other ACP expressed by *Scalindua* ^10^.

Close to this cluster of negatively charged residues on *amx*ACP, the Ser41 residue binds the phosphate group of the PPant-acyl moiety which extends away from *amx*ACP towards the *amx*FabF^het^ dimer. There, the PPant moiety, of which two conformations are observed for the part closest to *amx*ACP, is bound in the tunnel leading to the active site on *amx*FabF2, forming hydrogen bonds through Thr300 and Thr302. These hydrogen bonds are conserved in both the Iga11–Iga12 PKS/CLF complex ^32^ and in the *E. coli* AcpP–FabB complex ^44^, suggesting a preserved mechanism of PPant linker recognition across systems^32^. The catalytic triad binds the probe’s acyl group, with His298 and His335 forming an oxyanion hole binding the acyl oxygen, whereas the catalytic Cys165 binds covalently to the α-carbon of the C_8_ acyl unit. However, the electron density map for the alkyl chain breaks down as the substrate extends from Cys165, suggesting that the alkyl chain is disordered. Importantly, Cys165 was found to assume two conformations, only one of which is commensurate with probe binding. Likewise, the so-called “Gate 1 loop”, which contains the “gating residue” Phe394, was found in two conformations (Figure 6b). As expected from its proposed function, one of the conformations of the “gating” residue Phe394 blocks the is access to the hydrophobic part of the tunnel system where the acyl chain of the substrate is believed to bind.

**Figure 6.**
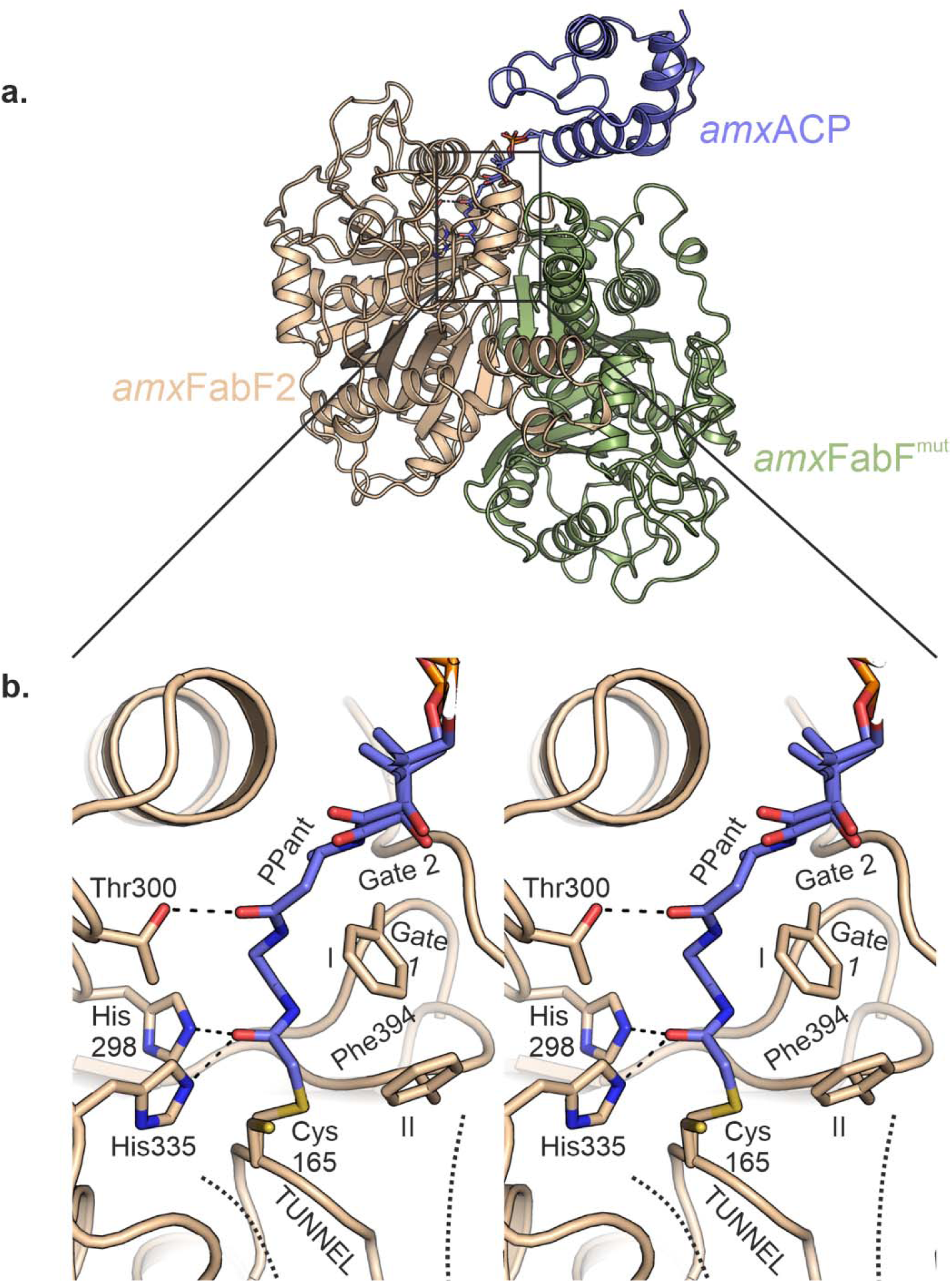
a. Overview of the *amx*ACP-*amx*FabF^het^ complex. The probe-labeled amxACP is shown in blue, *amx*FabF2 in beige and *amx*FabF^mut^ in green. **b.** Stereofigure, showing a closeup of the active site. The PPant-linker is hydrogen bonded to the conserved Thr300. The probe’s acyl oxygen is bound in the catalytic oxyanion hole formed by His298 and His355 from the catalytic triad, which is completed by Cys165. Cys165, in turn, is covalently bound to the probe’s C2 atom. Beyond this atom, the probe’s alkyl chain was not modeled for lack of electron density. Moreover, two conformations were observed for Cys165 as well as for the “Gate 1” loop containing the gating residue Phe394 (indicated as “I” and “II”); one of these is incompatible with the presence of probe and PPant linker. For the “Gate 2” loop, only one conformation was observed (see text). The continuation of the putative substrate binding tunnel further into the protein is indicated with dashed lines.

## Conclusions

The study of the mechanism of ladderane biosynthesis is complicated by the fact that so far, to our knowledge, no ladderane biosynthesis has been detected in *in* vitro settings, and that there is no genetic access to any ladderane-producing organism. So far, what little progress has been made was limited to bioinformatics analyses and labeling studies ^8, 14, 45^ as well as biochemical and structural studies on individual proteins from Cluster I ^9, 10^ though several mechanisms for ladderane biosynthesis have been proposed and a general consensus has emerged that Cluster I encodes the enzymes central to ladderane production. Indeed, it contains several of the enzymes that would be needed for fatty acid synthesis such as an acyl carrier protein (*amx*ACP) and a dehydratase (*amx*FabZ). Puzzlingly however, it contains two homologs of FabF, *amx*FabF2 and *amx*FabF^mut^. Based on sequence analysis, Rattray and coworkers had suggested that *amx*FabF2, with its functional catalytic triad, could catalyze chain elongation as in canonical fatty acid biosynthesis, whereas the noncanonical FabF homolog *amx*FabF^mut^ might catalyze another reaction, such as for instance the coupling between a short linear fatty acid and a separately produced ladderane moiety ^8^. However, while there are FabF homologs capable of elongating acyl chains by other units than a malonate-derived C_2_ moiety, a catalytic role for *amx*FabFmut seems unlikely given that of the catalytic triad the cysteine has mutated to a valine or threonine and the two histidines to glutamate, cysteine or asparagine, depending on the organism. These mutations would completely abrogate any ketosynthase activity.

Rather, the sequence- and biochemical analyses, crystal structures, and binding studies presented here paint a different picture: rather than functioning as two separate proteins, *amx*FabF2 and *amx*FabF^mut^ form a heterodimeric complex we call *amx*FabF^het^. Moreover, the data suggest that, in analogy to ketosynthase/chain length factor complexes, *amx*FabF^mut^ affects the chain length preference of the complex. This is the first time a chain length factor has been found in a fatty acid biosynthesis pathway.

While it is difficult to assess the substrate preference the complex shows when catalyzing its likely physiological reaction (chain elongation) from the results obtained using the crosslinking reaction employed here, which is mechanistically quite different, the complex would appear to favor the C_6_ and C_8_ substrates over long C_16_ substrates during the crosslinking reaction at least. This is in line with results obtained for the dehydratase *amx*FabZ, which was found to prefer short substrates during *in vitro* dehydration reactions ^10^. As discussed for *amx*FabZ, a preference for short chains for *amx*FabF and *amx*FabZ might indicate that these enzymes are involved in producing a short linear part of a typical ladderane lipid (Fig. 1a) which is typically 6 or 8 carbon atoms long. Experimental validation of this hypothesis would, however, require *in vitro* reconstitution of the entire ladderane biosynthetic pathway or genetic access to a ladderane-producing organism.

## Acknowledgements

We thank the staff of the Swiss Light Source (Villigen CH) for their support and facilities, and Chris Roome for outstanding computing support. We are also greatly indebted to Gunter Stier and Mike Jetten for the gift of genetic material, and to Ilme Schlichting for expert help with crystallization and data collection. Furthermore, T.R.M.B. is very grateful to Ilme Schlichting for her continuous support over many years. This work was funded by the Max Planck Society as well as by ERC Consolidator Grant 724362-STePLADDER to T.R.M.B.

