## Supplemental Information for "β-Ketoacyl Synthase II Homologs from a Ladderane-Producing Organism Form a Ketosynthase/Chain Length Factor-like functional heterodimer"

**Table of contents:**

**1. Biological and biochemical protocols**

1A. Production of 4’-phosphopantetheinyl transferase Sfp

1B. Production of CoA Biosynthesis Enzymes

**2. Chemical syntheses**

2A. Synthesis of hexanoyl-CoA thioester for MS-based activity assay

2B. General protocol for the synthesis of 2-bromo-acyl-phosphopantetheinamide linkers

**3. Supplementary Figures and Tables**

Supplementary Figure S1. Multiple sequence alignment

Supplementary Figure S2. Heterodimer between *amx*FabF2 and *amx*$FabF^{mut}$

Supplementary Figure S3. FabF dimer interfaces at the helix-turn-helix and interacting helix motifs

Supplementary Figure S4. Acyl-ACP probes for crosslinking assay

Supplementary Figure S5. Schematic of the mechanism-based crosslinking assay

Supplementary Figure S6. Enzymatic synthesis of the α-bromoacyl-ACP crosslinking probes

Supplementary Figure S7. Time-Course Crosslinking Assays with $C_6$-ACP Probe

Supplementary Figure S8. Time-Course Crosslinking Assays with $C_8$-ACP Probe

Supplementary Figure S9. Time-Course Crosslinking Assays with $C_{16}$-ACP Probe

Supplementary Figure S10. Interaction predictions with MCMAP

Supplementary Table 1. Composition of purification buffers for *B. fulgida* and *S. brodae* Constructs

Supplementary Table 2. Data collection and refinement statistics

### Additional materials and methods

#### 1. Biological and biochemical protocols

#### 1A. Production of 4’-phosphopantetheinyl transferase Sfp

The vector pET29b-BsSfp, encoding the *Bacillus subtilis* 4′-phosphopantetheinyl transferase (Sfp) with a C-terminal His-tag, was a gift from Michael Burkart (Addgene plasmid #75015; https://www.addgene.org/75015).

Chemically competent *E. coli* BL21 (DE3) cells were transformed with pET29b-BsSfp. A single colony was inoculated into 100 mL of Lysogeny Broth (LB) supplemented with 50 µg/mL kanamycin and cultured at 37 °C until the OD600 reached 1.5. This preculture (20 mL) was used to inoculate 2 L of LB medium containing 50 µg/mL kanamycin in a 5 L baffled Erlenmeyer flask. Cells were incubated at 37 °C with shaking at 100 rpm. Upon reaching an OD600 of 1.6, the temperature was lowered to 20 °C, and expression was induced with 500 µM isopropyl β-D-1-thiogalactopyranoside (IPTG). The culture was incubated overnight with shaking at 90 rpm.

Cells were harvested by centrifugation (6,000×g, 15 min, 4 °C) and resuspended in Wash Buffer (20 mM Tris/HCl pH 8.0, 500 mM NaCl, 10 mM imidazole), supplemented with hen egg-white lysozyme (Sigma-Aldrich) and DNaseI (Roche). Cell lysis was performed by passing the suspension five times through a Microfluidizer (Microfluidics, USA). The lysate was cleared by centrifugation at 50,000×g for 1 h at 4 °C and filtered through a 0.45 µm syringe filter.

The lysate was loaded onto a 5 mL Ni-NTA agarose gravity-flow column (Qiagen) equilibrated with Wash Buffer. The column was washed with 30 mL of Wash Buffer, and the protein was eluted using Wash Buffer supplemented with 300 mM imidazole. Fractions containing Sfp were pooled and mixed with TEV protease (16 µM) and DTE (5 mM). The mixture (10 mL) was dialyzed overnight at 4 °C against Dialysis Buffer (20 mM Tris/HCl pH 8.0, 150 mM NaCl, 2 mM DTE, 5% (v/v) glycerol).

To remove the cleaved His-tag and TEV protease, the dialysate was passed twice through a 5 mL Ni-NTA agarose column. The flow-through and wash fractions containing the cleaved Sfp were dialyzed twice against Storage Buffer (10 mM Tris/HCl pH 7.5, 1 mM EDTA, 10% (v/v) glycerol) at 8 °C. The purified protein was concentrated using 10 kDa MWCO Amicon concentrators (Millipore) to a final volume of approximately 3 mL. Protein purity was assessed by 15% SDS-PAGE. The final concentration was determined to be 29 mg/mL (1.08 mM) based on absorbance at 280 nm ($\epsilon_{280}$=30,370 $M^{-1}cm^{-1}$). Aliquots (100 µL) were flash-frozen in liquid nitrogen and stored at −80 °C.

**1B. Production of CoA Biosynthesis Enzymes**

The plasmids pET30_PanK (C-His), pET39_PPAT (C-His), and pET28a_DPCK (C-His), encoding the *E. coli* CoA biosynthesis enzymes pantothenate kinase (PanK), phosphate adenyltransferase (PPAT), and dephospho-CoA kinase (DPCK), respectively, were generously provided by T. Kittilä and M. Cryle (MPI for Medical Research). Protein production was performed as described previously [1].

Transformed *E. coli* BL21(DE3) cells were cultured in 4 L of LB medium (two 2 L cultures) supplemented with 50 µg/mL kanamycin at 37 °C with shaking at 100 rpm. Protein expression was induced by adding 500 µM IPTG when the culture reached an $OD_{600}$ of approximately 0.6. The temperature was then reduced to 20 °C, and the culture was incubated overnight with shaking at 90 rpm.

*Lysis and Ni-NTA Affinity Chromatography*. Cells were harvested by centrifugation at 6,000×g for 15 min at 4 °C. The resulting cell pellet was resuspended in Washing Buffer (WB; 150 mM NaCl, 50mM Tris-HCl, pH 8.0, 10 mM imidazole). Cell disruption was achieved by sonication using a probe sonicator (3 cycles of 1 min total sonication time with 0.5 s on/0.5 s off pulses at 50%

amplitude). Cell debris and insoluble protein were removed by centrifugation at 20,000 rpm (47,800×g equivalent) for 40 min at 4 °C.

The cleared lysate was sterile-filtered (0.45 μm) and loaded onto a 5 mL Ni-NTA gravity flow column, which had been pre-equilibrated with WB. Bound protein was washed with 5 column volumes of WB. The protein was eluted with Elution Buffer (EB; 150 mM NaCl, 50 mM Tris-HCl, pH 8.0, 300 mM imidazole).

*Dialysis and Size Exclusion Chromatography (SEC).* Pooled protein-containing fractions were dialyzed overnight against Dialysis Buffer (150 mM NaCl, 50 mM Tris-HCl, pH 8.0, 2 mM $MgCl_2$) and then concentrated using 10 kDa MWCO (molecular weight cut-off) Amicon filters. The proteins were further purified by SEC using a Superdex 75 (16/60) column connected to an ÄKTA PURE system, employing the Dialysis Buffer as the running buffer. Protein purity was assessed by 15% SDS-PAGE. The purified proteins were aliquoted, flash-frozen in liquid nitrogen, and stored at −80 °C.

### 2. Chemical Syntheses

Chemicals were analysis grade and used without further purification. Thin layer chromatography (TLC) was performed on either normal-phase Polygram SIL G/UV 254 silica plates or reverse-phase RP-18 W/UV254 plates (obtained from Macherey Nagel GmbH, Düren, Germany). Flash chromatography was performed on an Isolera One System (Biotage AB, Uppsala, Sweden) using either KP-SIL (Biotage AB, Uppsala, Sweden) or SiliaSep (SiliCycle Inc., Quebec City, Canada) normal-phase silica columns. Atmospheric Pressure Chemical Ionization (APCI) or Electrospray Ionization (ESI) mass spectra were recorded using an Expression L compact mass spectrometer equipped with a Plate Express TLC extraction device (Advion Inc., Ithaca, NY, USA). $^{1}H$ and $^{13}C$ nuclear magnetic resonance (NMR) spectra were measured on a Bruker Avance III 400 MHz spectrometer equipped with a cryoprobe. NMR spectra were referenced to the indicated residual deuterated solvent signal. Chemical shifts δ are given in ppm, coupling constants *J* in Hertz (Hz). Multiplicities are abbreviated as follows: s = singlet, d = doublet, t = triplet, q = quartet, p = quintet, h = sextet, m = multiplet and bs = broad signal. NMR data were processed using MestReNova v11.0.4 (Mestrelab Research, Santiago de Compostela, Spain).

### 2A. Synthesis of hexanoyl-CoA thioester for MS-based activity assay

Acyl-CoA thioesters were synthesized for use as primers in the *in vitro* reconstitution and MS-based assays. The synthesis of hexanoyl-CoA was performed via a two-step procedure: the initial formation of hexanoic acid N-hydroxysuccinimide (NHS) ester, followed by its reaction with coenzyme A (CoA-SH), adapted from Mukherjee *et al.* [2].

*Synthesis of 2,5-dioxopyrrolidin-1-yl hexanoate (1)*

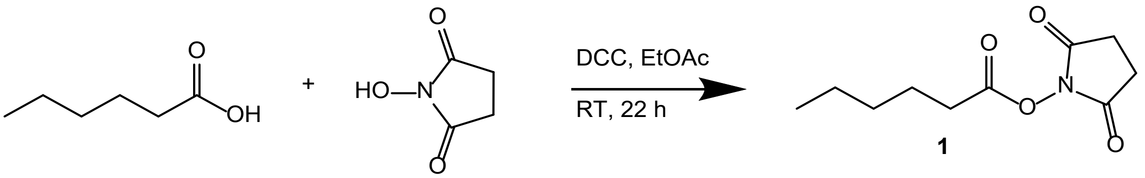


A round-bottom flask was charged with hexanoic acid (1.20 mmol, 1 eq) and N-Hydroxysuccinimide (1.20 mmol, 1 eq), dissolved in 40 mL of ethyl acetate (EtOAc). A solution of N,N′-dicyclohexylcarbodiimide (DCC; 1.20 mmol, 1 eq) in 10 mL of EtOAc was added, and the reaction mixture was stirred for 22 hours at room temperature. The resulting white precipitate of dicyclohexylurea (DCU) was removed by filtration through Celite. The filtrate was evaporated *in vacuo*, and the crude product was purified by automated flash column chromatography (Isolera system, 25 g HP silica cartridge). Purification was performed using a linear gradient of 20% to 100% ethyl acetate in hexane. Fractions containing the pure product were pooled, and the solvent was removed *in vacuo* to yield 182 mg (71 % yield) of hexanoyl-NHS ester **1** as a colorless solid.

$^1$H NMR (400 MHz, $CDCl_3$): δ (ppm) = 0.91 (t, J=7.0, 3H, C$H_3^{1}$), 1.27 – 1.46 (m, 4H, C$H_2^{2,3}$), 1.74 (p, J=7.5, 2H, C$H_2^{4}$), 2.59 (t, J=7.5, 2H, C$H_2^{5}$), 2.83 (s, 4H, C$H_2^{12,13}$).

$^{13}$C NMR (101 MHz, $CDCl_3$): 13.94 ($C^{1}H_3$), 22.28 ($C^{2}H_2$), 24.38 ($C^{4}H_2$), 25.73 ($C^{12,13}H_2$), 31.04 ($C^{3,5}H_2$), 168.83 ($C^{6}O$), 169.33 ($C^{10,14}O$).

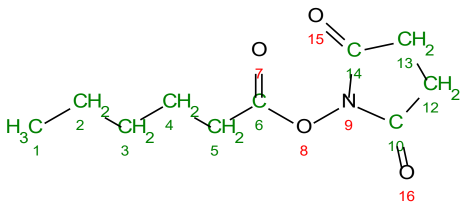

MS: $C_{10}H_{15}NO_4$ exact 213.1001, $[M+H]^+$ calculated 214.1, found 214.1 APCI(+)

*Synthesis of Hexanoyl-CoA (2)*

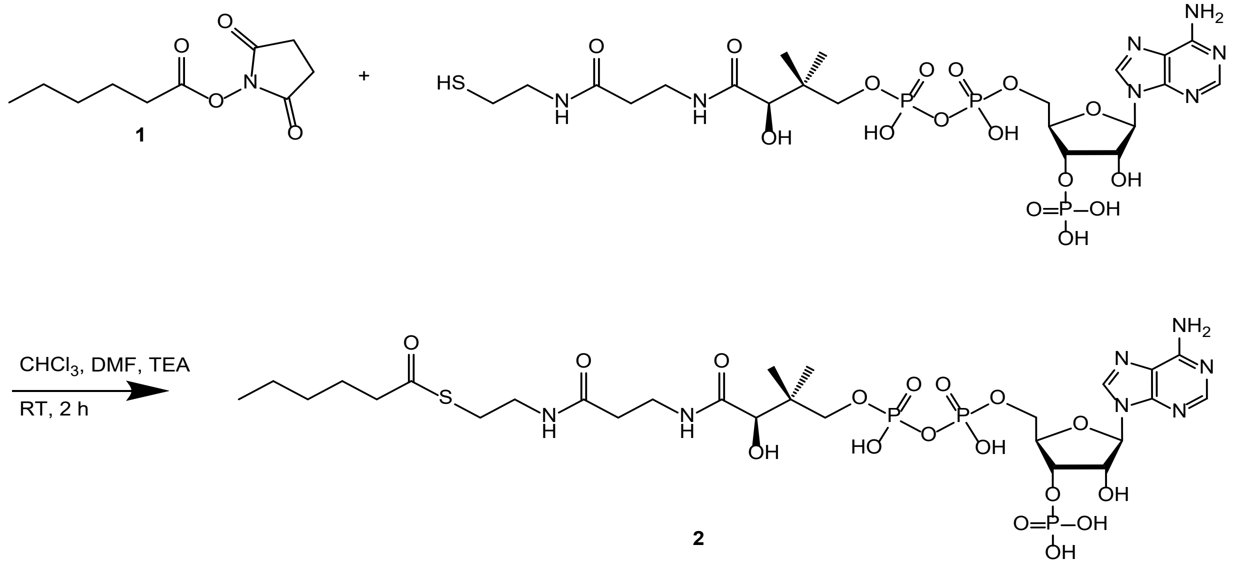


Coenzyme A (free acid form, 6.09 µmol, 1 eq) was suspended in dry chloroform ($CHCl_3$) and solubilized by the addition of 2 M triethylamine (TEA) solution in $CHCl_3$ (4 eq). The hexanoyl-NHS ester **1** (9.13 µmol, 1.5 eq), dissolved in 46 µL of a 4:1 (v/v) mixture of dry $CHCl_3$ and dimethyl formamide (DMF), was added to the CoA solution. The reaction was allowed to proceed for 2 hours at room temperature with mixing.

The resulting product was washed three times with diethyl ether ($Et_2O$) to remove unreacted NHS ester and other impurities. The final residue was dried, redissolved in 2% (v/v) aqueous formic acid, and purified by Solid-Phase Extraction (SPE) on a 1 mL C18 column. The column was conditioned with methanol and water, and equilibrated with 2% (v/v) formic acid in water. After loading the sample, the column was washed with 2% formic acid, and the final hexanoyl-CoA product was eluted with methanol. The purity of the fractions was confirmed by RP-TLC-ESI-MS. After evaporation of the solvent 27 mg (crude/wet, 116 % apparent yield) of compound **2** were obtained.

MS: $C_{27}H_{46}N_7O_{17}P_3S$ exact 865.1884, $[M-2H]^{2-}$ calculated m/z 431.6, found 431.7 ESI(-)

**2B. Synthetic Methods for Phosphopantetheine and α-Bromoacyl Derivatives (Linkers)**

Syntheses were performed using methods described in [3-5].

*Synthesis of PMP-protected (R)-pantothenic acid (3)* [3]

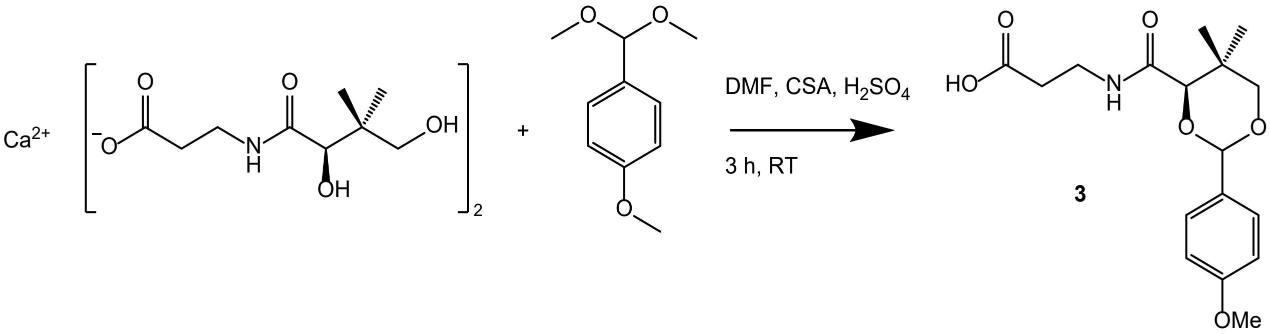


D-Pantothenic acid hemicalcium salt (47.65 g, 100 mmol) was dissolved in 900 mL of DMF in a 2 L round-bottom flask, fitted with a calcium chloride drying tube. To the solution, 40 g of 4A molecular sieves was added to absorb water and methanol. Concentrated sulfuric acid (5.6 mL, 100 mmol) was then added dropwise while stirring continuously. The reaction mixture was stirred for 1 hour, forming a gel-like, brownish slurry, with the partial dissolution of the molecular sieve. Subsequently, *p*-anisaldehyde dimethyl acetal (34.1 mL, 200 mmol) and camphorsulfonic acid (CSA, 4.65 g, 20 mmol) were introduced to the mixture. The resulting brownish slurry was stirred at room temperature for 2 hours. Following this reaction period, 1 L of water was added to the reaction mixture, which was then vacuum filtered through a Celite pad, with additional water used to wash the filtrate. The yellowish filtrate (total volume 1200 mL) was divided into two equal parts (600 mL each). Each portion was extracted three times with 400 mL of ethyl acetate. The combined organic extracts (2400 mL) were divided into two portions of 1200 mL each, which were then washed three times with 200 mL of water. The organic phases were dried over anhydrous sodium sulfate ($Na_2SO_4$), and ethyl acetate was removed under reduced pressure (< 25 mbar, 15 min) using a rotary evaporator to yield a yellow/white viscous syrup (suspension) weighing 45.41 g. This crude product was dried using a lyophilizer for 15 hours, resulting in 35.88 g of solid. The dried product was resuspended in 80 mL of cold dichloromethane (DCM) and filtered under vacuum, followed by washing with 20 mL of DCM to remove any residual *p*-anisaldehyde dimethyl acetal. This process yielded 11.08 g of the desired product as a white crystalline solid (Fraction A, 89 % purity). The remaining DCM in the filtrate was removed by rotary evaporation under reduced pressure (< 20 mbar, 20 min), resulting in 24.07 g of product. This product was further dried using a lyophilizer for 40 hours to yield 21.13 g of a yellow syrup. The syrup was resuspended in 20 mL of cold DCM and 10 mL of hexane, then filtered under vacuum and washed with 5 mL of DCM, resulting in 2.01 g of a white solid (Fraction B, 79% purity). The total yield of compound **3** was about 17%.

$^{1}$H NMR (400 MHz, $CDCl_3$): δ (ppm) = 1.09 (s, 3H, C$H_3^{12}$), 1.10 (s, 3H, C$H_3^{11}$), 2.61 (t, *J*=6.2, 2H, C$H_2^{5}$), 3.44 – 3.61 (m, 2H, C$H_2^{4}$), 3.68 (q, *J*=11.2, 2H, C$H_2^{1}$), 3.81 (s, 3H, C$H_3^{24}$), 4.10 (s, 1H, C$H^{10}$), 5.46 (s, 1H, C$H^{20}$), 6.88 – 6.94 (m, 2H, C$H^{15,17}$), 7.37 – 7.44 (m, 2H, C$H^{14,18}$).

$^{13}$C NMR (101 MHz, $CDCl_3$): 19.23 ($C^{11}H_3$), 21.94 ($C^{12}H_3$), 33.26 ($C^{9}$ *q*), 33.95 ($C^{5}H_2$), 34.24 ($C^{4}H_2$), 55.46 ($C^{24}H_3$), 78.62 ($C^{1}H_2$), 83.85 ($C^{10}H$), 101.37 ($C^{20}H$), 113.89 ($C^{15,17}H$), 127.57 ($C^{14,18}H_3$), 132.17 ($C^{19}q$), 160.35 ($C^{16}q$), 169.65 ($C^{3}O$ *q*), 176.58 ($C^{7}OOH$ *q*).

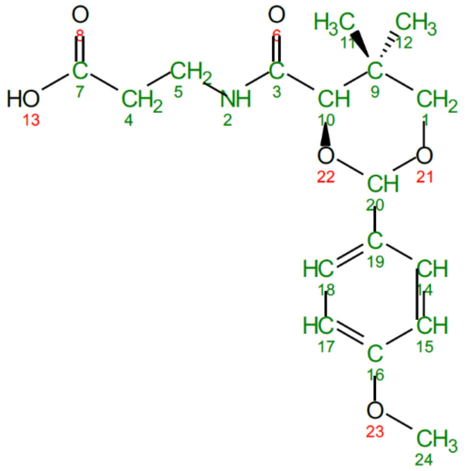

MS: $C_{17}H_{23}NO_6$ exact 337.1525, $[M+H]^+$ calculated 338.2, found 338.2 APCI (+)

*Synthesis of 2-azido-ethylamine (4)* [3]

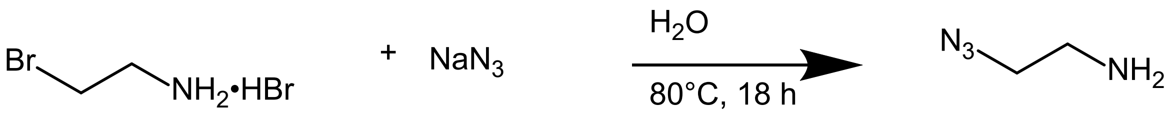


**4**

A 500 mL three-necked flask equipped with a Dimroth condenser and a thermometer was charged with 2-bromoethylamine hydrobromide (41.0 g, 200 mmol) dissolved in 200 mL of water. Sodium azide (39.0 g, 600 mmol) was added to the solution. The reaction mixture was stirred and heated at 80 °C in an oil bath maintained at 100 °C for 16 hours. The following day, the reaction mixture was cooled to room temperature and then further cooled to 0 °C using an ice-water bath. Solid potassium hydroxide (KOH, 60 g) was gradually added to the mixture and stirred until completely dissolved. Diethyl ether (200 mL) was then added, and the solution was stirred for 35 minutes. The organic phase was separated, and the aqueous layer was extracted three times with 400 mL of diethyl ether ($Et_2O$). The combined organic layers were dried over sodium sulfate ($Na_2SO_4$), and the solvent was removed under reduced pressure (minimum 200 mbar) to yield 11.82 g of a yellowish oil. The crude product was filtered through a layer of $Na_2SO_4$ using an Allihn tube, yielding 11.19 g (92% purity, 60% yield) of 2-azido-ethanamine **4**.

$^{1}$H NMR (400 MHz, $CDCl_3$): δ (ppm) =

1.40 (s, 2H, $NH_2^1$), 2.86 (t, J=5.7, 2H, C$H_2^3$), 3.35 (t, J=5.7, 2H, C$H_2^2$)

$^{13}$C NMR (101 MHz, $CDCl_3$): 41.46 (C $^2H_2$), 54.76 (C$^3H_2$)

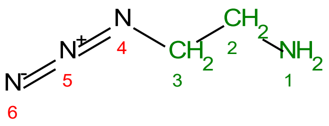


MS: $C_2H_6N_4$ exact 86.0592, $[M+H]^+$ calculated: 87.1; found 87.0 ESI (+)

*Synthesis of (4R)-N-(3-((2-azidoethyl) amino)-3-oxopropyl)-2-(4-methoxyphenyl)-5,5-dimethyl-1,3-dioxane-4-carboxamide (5)* [3, 4]

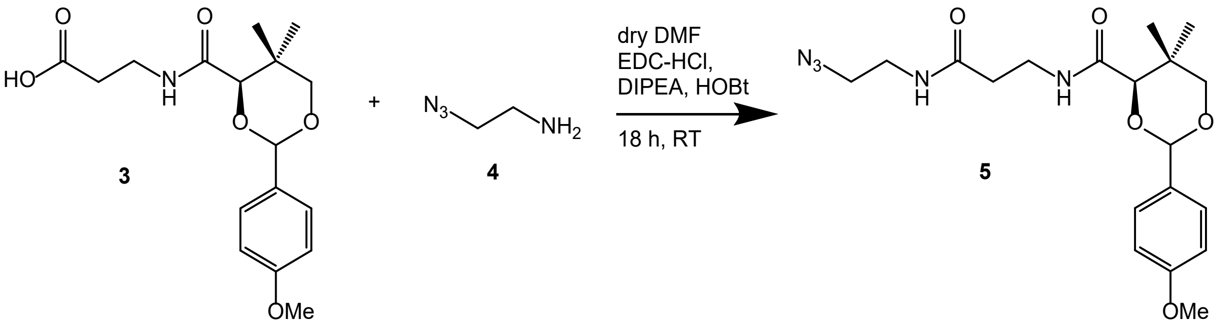


In a 100 mL flask equipped with a calcium chloride drying tube, PMP-protected pantothenate **3** (12.34 g, 32 mmol) was dissolved in 30 mL of dry DMF. To this solution, 2-azidoethanamine **4** (3.0 g, 32 mmol) and hydroxybenzotriazole (HOBt, 5.9 g, 38.4 mmol, 1.2 eq, hydrate ~12% $H_2O$) were added. The mixture was stirred with 10 g of molecular sieve 3A for 35 minutes, during which it turned yellow and warmed up. Subsequently, 1-Ethyl-3-(3-dimethylaminopropyl)carbodiimid hydrocloride (EDC-HCl, 7.36 g, 38.4 mmol, 1.2 eq) and Diisopropylethylamin (DIPEA, 14 mL, 80 mmol, 2.5 eq) were added, and the reaction was allowed to stir at room temperature overnight, resulting in a brown/red colored mixture. The reaction mixture was filtered through a thin layer of silica using pressure filtration in an Allihn tube and washed with 50 mL of ethyl acetate (EtOAc). The deep orange filtrate was stirred with 150 mL of water for 5 minutes and then extracted three times with 150 mL of EtOAc. The combined organic phases were washed 3 x with 100 mL of saturated sodium bicarbonate ($NaHCO_3$) solution and 3 x 100 mL of brine. The organic layer was dried over sodium sulfate ($Na_2SO_4$). The ethyl acetate extract (14.04 g) was loaded onto a short silica gel column (350 mL) and eluted with a gradient of hexane from 30:70 to 100% EtOAc. Fractions containing the expected product were pooled, and the solvent was evaporated under reduced pressure to yield a yellowish/orange viscous oil (13.021 g). This material was dried in a lyophilizer for 1 hour to yield 11.305 g of a yellowish resin (78% purity, 68% yield).

$^1H$ NMR (400 MHz, $CDCl_3$): δ (ppm) = 1.10 (s, 3H, C$H_3^{12}$), 1.11 (s, 3H, C$H_3^{11}$), 2.44-2.48 (m, 2H, C$H_2^{5}$), 3.37 – 3.39 (m, 2H, C$H_2^{23}$), 3.42 – 3.47 (m, 2H, C$H_2^{24}$), 3.52 – 3.58 (m, 2H, C$H_2^{4}$),

3.69 (q, $J$=11.7, 2H, C$H_2^{1}$), 3.82 (s, 3H, C$H_3^{27}$), 4.09 (s, 1H, C$H^{10}$), 5.46 (s, 1H, C$H^{20}$), 6.25 (bs, 1H, NH$^{13}$), 6.90 – 6.95 (m, 2H, C$H^{15,17}$), 7.37 – 7.47 (m, 2H, C$H^{14,18}$).

$^{13}$C NMR (101 MHz, $CDCl_3$): 19.26 (C$^{11}H_3$), 21.98 (C$^{12}H_3$), 33.25 (C$^{9}q$), 34.94 (C$^{4}H_2$), 36.28 (C$^{5}H_2$), 39.03 (C$^{24}H_2$), 50.85 (C$^{23}H_2$), 55.50 (C$^{27}H_3$), 78.63 (C$^{1}H_2$), 83.97 (C$^{10}H$), 101.53 (C$^{20}H$), 113.90 (C$^{15,17}H$), 127.66 (C$^{14,18}H$), 130.28 (C$^{19}q$), 160.42 (C$^{16}q$), 169.87 (C$^{3}O$), 171.29 (C$^{7}O$).

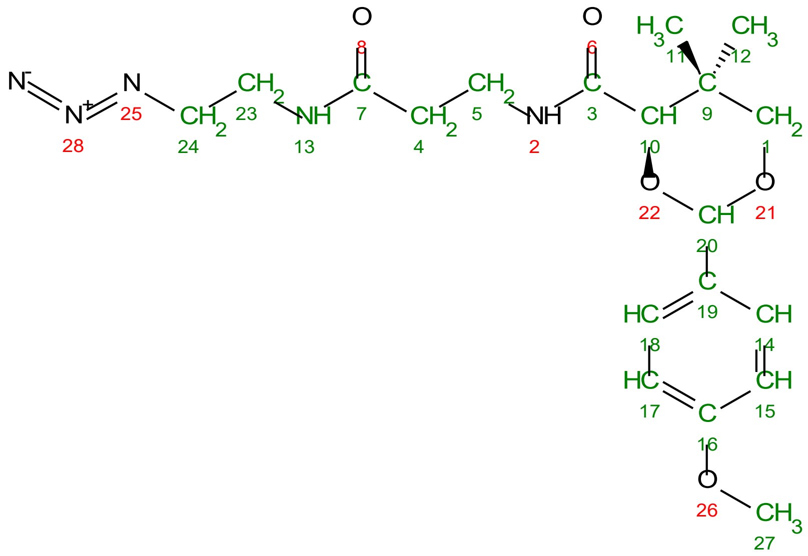

MS: $C_{19}H_{27}N_5O_5$ exact 405.2012, $[M+H]^+$ calculated 406.2, found 406.2 APCI (+), $[M-H]^-$ calculated 404.2, found 404.1 APCI (-)

*Synthesis of (4R)-N-(3-((2-aminoethyl)amino)-3-oxopropyl)-2-(4-methoxyphenyl)-5,5-dimethyl-1,3-dioxane-4-carboxamide (6)* [3, 4]

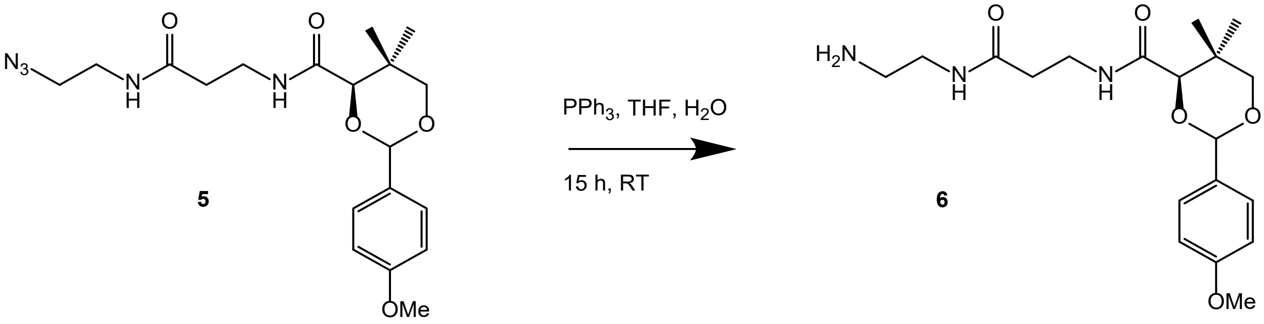


In a 50 mL flask, azide **5** (11.3 g, ~27 mmol, 1.0 eq) and triphenylphosphine ($PPh_3$, 7.79 g, 29.7 mmol, 1.1 eq) were dissolved in 22 mL of tetrahydrofuran (THF) and 2.2 mL of water (4.5 eq). The mixture was stirred at room temperature overnight (15 hours), during which $N_2$ bubbles formed initially. The reaction progress was monitored by thin-layer chromatography (TLC) using silica gel and a solvent system of DCM/MeOH + $Et_3N$ (85:15). Solvents were evaporated under reduced pressure (<20 mbar, 15 minutes), and the resulting amine **6** was isolated by flash chromatography. The elution was performed using a gradient from DCM + 2% $Et_3N$ to DCM/MeOH (85:15). The solvent from the collected fractions was removed by rotary evaporation, yielding 11.521 g of a viscous, colorless oil. The crude product was dissolved in 100 mL of DCM and washed twice with 100 mL of water and twice with 100 mL of brine. The aqueous layers were extracted twice with 50 mL of DCM. The combined organic layers were dried over $Na_2SO_4$, and the solvent was removed by rotary evaporation (<15 mbar, 45 minutes), yielding 8.991 g of the product. The product was then dried on a lyophilizer for 17 hours, resulting in 8.753 g (85%) of a colorless, viscous, fluffy powder, that was dissolved in THF (380 mg/mL), filtered through a layer of $Na_2SO_4$, and stored at 8 °C.

$^1$H NMR (400 MHz, $CDCl_3$): δ (ppm) = 1.08 (s, 3H, C$H_3^{12}$), 1.10 (s, 3H, C$H_3^{11}$), 2.44 (t, *J*=6.3, 2H, C$H_2^{5}$), 2.85 (t, *J*=5.7, 2H, C$H_2^{24}$), 3.27 – 3.39 (m, 3H, C$H_2^{23}$), 3.51 – 3.59 (m, 2H, C$H_2^{4}$), 3.69 (q, *J*=11.2, 2H, C$H_2^{1}$), 3.82 (s, 3H, C$H_3^{27}$), 4.08 (s, 1H, C$H^{10}$), 5.46 (s, 1H, C$H^{20}$), 6.62 (t, *J*=5.7, 1H, $NH^{13}$), 6.88 – 6.96 (m, 2H, C$H^{15,17}$), 7.05 (t, *J*=6.3, 1H, $NH^{2}$), 7.39 – 7.49 (m, 2H, C$H^{14,18}$).

$^{13}C$ NMR (101 MHz, $CDCl_3$): 19.30 ($C^{11}H_3$), 22.03 ($C^{12}H_3$), 33.24 ($C^{9}q$), 35.27 ($C^{4}H_2$), 36.41 ($C^{5}H_2$), 41.06 ($C^{23,24}H_2$), 55.50 ($C^{27}H_3$), 78.62 ($C^{1}H_2$), 84.00 ($C^{10}H$), 101.51 ($C^{20}H$), 113.90 ($C^{15,17}H$), 127.68 ($C^{14,18}H$), 130.29 ($C^{19}q$), 160.41 ($C^{16}q$), 169.79 ($C^{3}O$), 171.55 ($C^{7}O$).

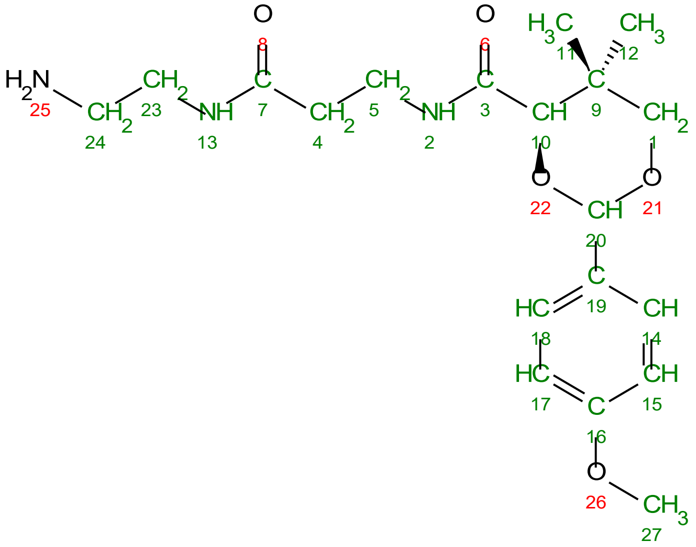

MS: $C_{19}H_{29}N_3O_5$ exact 379.2107, $[M-H]^-$ calculated 378.2, found 378.1 APCI (-)

*General protocol for the synthesis of 2-bromo-acyl-phosphopantetheinamide linkers (7a-c)*

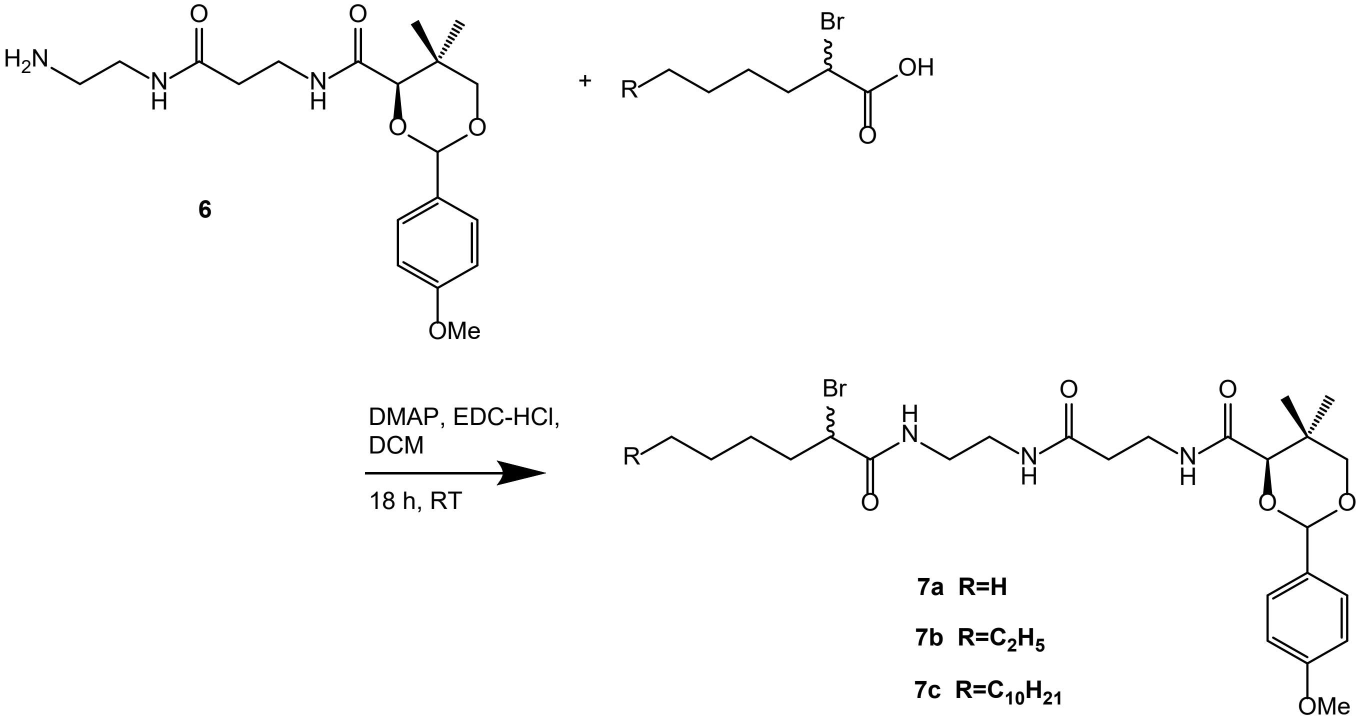


In a 10 mL pear-shaped flask, (4R)-N-(3-((2-aminoethyl)amino)-3-oxopropyl)-2-(4-methoxyphenyl)-5,5-dimethyl-1,3-dioxane-4-carboxamide **6** (266 µmol, 1.0 eq), 2-bromo-acyl acid (399 µmol, 1.5 eq), and 2.7 mL of dichloromethane ($CH_2Cl_2$) were combined. To this mixture, 1-Ethyl-3-(3-dimethylaminopropyl)carbodiimid hydrocloride (EDC-HCl, 444 µmol, 1.7 eq) and 4-(Dimethylamino)pyridin (DMAP, 79.4 µmol, 0.30 eq) were added. The reaction was allowed to proceed overnight. Subsequently, the reaction mixture was diluted with 100 mL of $CH_2Cl_2$ and washed with 10 mL of brine. The organic phase was dried over anhydrous sodium sulfate ($Na_2SO_4$), filtered, and concentrated under reduced pressure using rotary evaporation. The crude products were purified by silica gel (30 g) flash chromatography using a 39:1 mixture of dichloromethane and methanol.

*(4R)-N-(3-((2-(2-bromo-hexyl)ethyl)amino)-3-oxopropyl)-2-(4-methoxyphenyl)-5,5-dimethyl-1,3-dioxane-4-carboxamide (7a)*

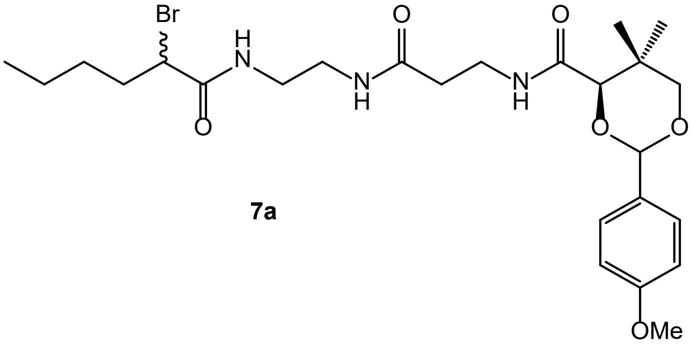


**7a**

Starting from 2-bromo-hexanoic acid, white solid used without determining yield.

MS: $C_{25}H_{38}BrN_3O_6$ exact 555.1944, [M+H]+ calculated 556.2, found 556.3 APCI(+)

*(4R)-N-(3-((2-(2-bromo-octyl)ethyl)amino)-3-oxopropyl)-2-(4-methoxyphenyl)-5,5-dimethyl-1,3-dioxane-4-carboxamide (7b)*

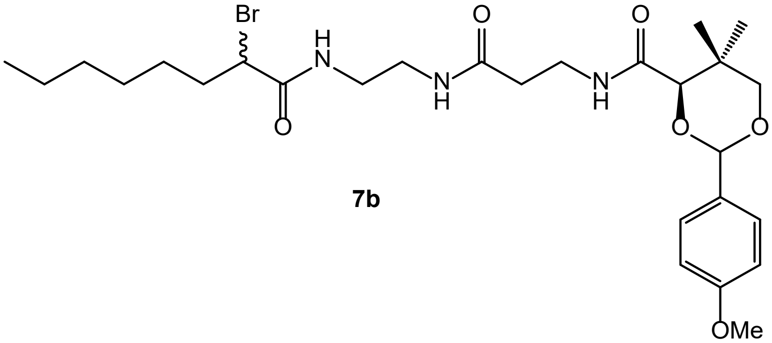


**7b**

Starting from 2-bromo-octanoic acid, white solid (90 mg, 58% yield)

MS: $C_{27}H_{42}BrN_3O_6$ exact 583.2257, $[M+H]^+$ calculated 584.2, found 585.2 APCI (+)

*(4R)-N-(3-((2-(2-bromo-hexadecyl)ethyl)amino)-3-oxopropyl)-2-(4-methoxyphenyl)-5,5-dimethyl-1,3-dioxane-4-carboxamide (7c)*

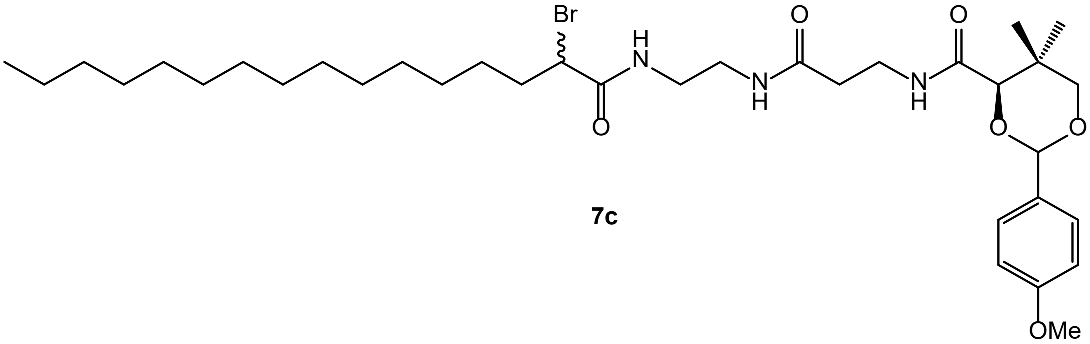


Starting from 2-bromo-hexadecanoic acid, white solid (172 mg, 77 % yield)

MS: $C_{35}H_{58}BrN_3O_6$ exact 695.3509, $[M+H]^+$ calculated 696.4, found 696.8 APCI (+)

*Deprotection of 2-bromo-acyl-phosphopantetheinamide linkers (8a-c)*

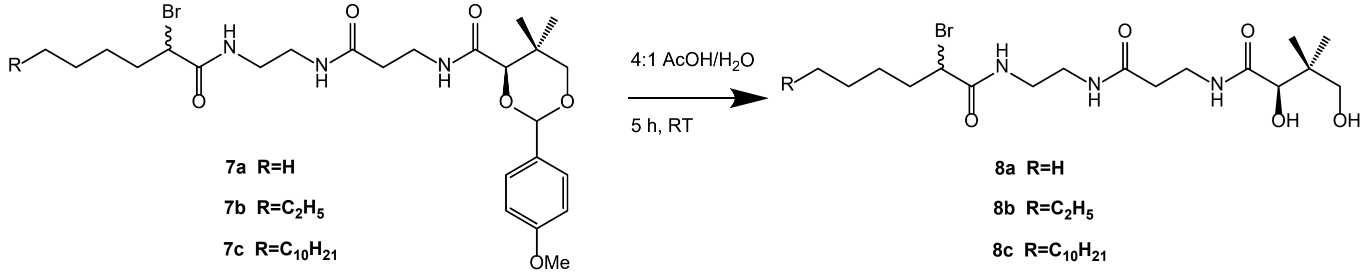


In a 20 mL vial, compounds **7a-c** (220 µmol, 1.0 eq.) were combined with 2.2 mL of a 4:1 mixture of acetic acid (AcOH) and water ($H_2O$). The reaction mixture was allowed to proceed for 5 hours at room temperature, after which it was concentrated by rotary evaporation. The residue was subjected to azeotropic distillation using cyclohexane (3 × 10 mL) and toluene (3 × 10 mL). The crude product was then purified by silica gel flash chromatography, employing a gradient of dichloromethane/methanol (19:1 → 4:1).

*2-bromo-N-(2-(3-((R)-2,4-dihydroxy-3,3-dimethylbutanamido)propanamido)ethyl)hexanamide (8a)*

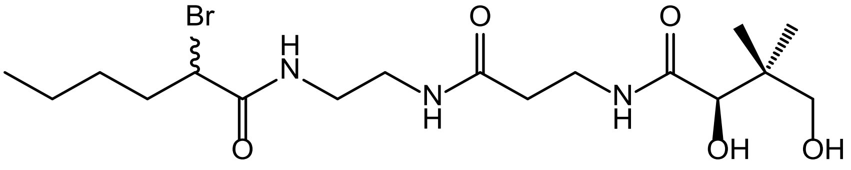


**8a**

Starting from **7a**, White solid (61 mg)

$^{1}$H NMR (400 MHz, MeOD): δ (ppm) = 0.92 (s, 3H, C$H_3^{25}$), 0.94 (s, 6H, C$H_3^{11,12}$), 1.31 – 1.43 (m, 2H, C$H_2^{23}$), 1.43 – 1.54 (m, 2H, C$H_2^{24}$), 1.90 – 2.08 (m, 2H, C$H_2^{19}$), 2.44 (t, J=6.9, 2H, C$H_2^{5}$), 3.17 – 3.36 (m, 4H, C$H_2^{16,17}$), 3.37 – 3.53 (m, 2H, C$H_2^{1}$), 3.45 – 3.53 (m, 2H, C$H_2^{4}$), 3.90 (s, 1H, C$H^{10}$), 4.29 (t, J=7.3, 1H, C$H^{20}$).

$^{13}$C NMR (101 MHz, MeOD): 14.20 (C$^{25}H_3$), 20.90 (C$^{12}H_3$), 21.35 (C$^{11}H_3$), 23.06 (C$^{23}H_2$), 23.40 (C$^{24}H_2$), 36.04 (C$^{19}H_2$), 36.33 (C$^{4}H_2$), 36.62 (C$^{5}H_2$), 40.19 (C$^{16,17}H_2$), 40.34 (C$^{9}q$), 49.71 (C$^{20}H$), 70.32 (C$^{1}H_2$), 77.33 (C$^{10}H$), 172.31 (C$^{21}O$), 174.15 (C$^{7}O$), 176.03 (C$^{3}O$).

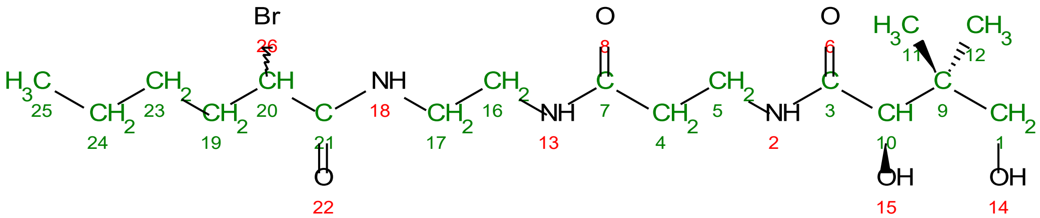


MS: $C_{17}H_{32}BrN_3O_5$ exact 437.1525, $[M+H]^+$ calculated 438.2, found 438.3 APCI(+)

*2-bromo-N-(2-(3-((R)-2,4-dihydroxy-3,3-dimethylbutanamido)propanamido)ethyl)octanamide (8b)*

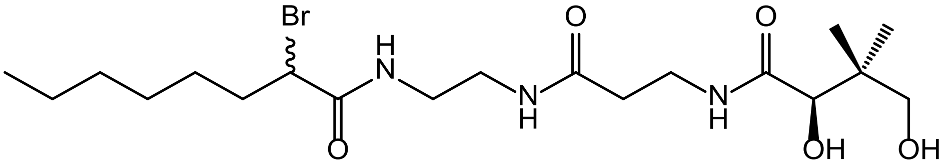


**8b**

Starting from **7b**, White solid (64 mg, 89 % yield)

$^{1}$H NMR (400 MHz, CDCl$_3$): δ (ppm) = 0.88 (t, J=6.6, 3H, C$H_3^{27}$), 0.94 (s, 3H, C$H_3^{11}$), 1.01 (s, 3H, C$H_3^{12}$), 1.24 – 1.35 (m, 6H, C$H_2^{24,25,26}$), 1.36 – 1.53 (m, 2H, C$H_2^{23}$), 1.90 – 2.13 (m, 2H, C$H_2^{19}$), 2.41 – 2.52 (m, 2H, C$H_2^{4}$), 3.26 – 3.40 (m, 2H, C$H_2^{5}$), 3.41– 3.57 (m, 4H, C$H_2^{16,17}$), 3.60 – 3.73 (m, 2H, C$H_2^{1}$), 4.03 (d, J=2.7, 1H, C$H^{10}$), 4.24-4.30 (m, 1H, C$H^{20}$).

$^{13}$C NMR (101 MHz, CDCl$_3$): 14.01 (C$^{27}H_3$), 20.59 (C$^{12}H_3$), 21.43 (C$^{11}H_3$), 22.51 (C$^{26}H_2$), 27.26 (C$^{23}H_2$), 28.48 (C$^{24}H_2$), 31.52 (C$^{25}H_2$), 35.34 (C$^{4}H_2$), 35.48 (C$^{5}H_2$), 36.09 (C$^{19}H_2$), 39.25 (C$^{9}q$), 39.27 (C$^{16}H_2$), 40.08 (C$^{17}H_2$), 50.73 (C$^{20}H$), 70.81 (C$^{1}H_2$), 77.20 (C$^{10}H$), 170.53 (C$^{21}O$), 172.38 (C$^{7}O$), 174.14 (C$^{3}O$).

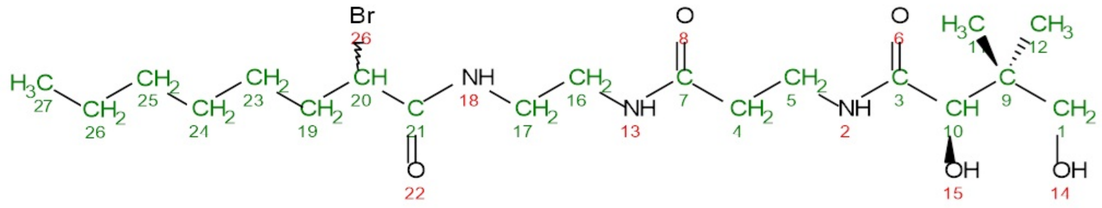

MS: $C_{19}H_{36}BrN_3O_5$ exact 465.1838, $[M+H]^+$ calculated 466.2, found 467.0 APCI(+)

*2-bromo-N-(2-(3-((R)-2,4-dihydroxy-3,3-dimethylbutanamido)propanamido)ethyl)*

*hexadecylamide (8c)*

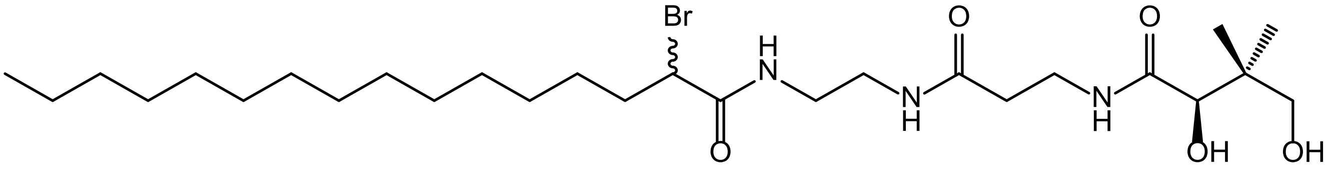

**8c**

Starting from **7c**, Yellowish solid (84 mg, 74 % yield)

$^{1}$H NMR (400 MHz, MeOD): δ (ppm) = 0.87 – 0.97 (m, 9H, C$H_3^{11,12,35}$), 1.30 (bs, 27H, $CH_2$), 1.59 – 1.74 (m, 2H, C$H_2^{34}$), 1.91 – 2.15 (m, 2H, C$H_2^{33}$), 2.34 – 2.49 (m, 2H, C$H_2^{5}$), 3.17 – 3.35 (m, 2H, C$H_2^{16,17}$), 3.34 – 3.54 (m, 2H, C$H_2^{1}$), 3.45 – 3.56 (m, 2H, C$H_2^{4}$), 3.91 (s, 1H, C$H^{10}$), 4.29 (t, $J$=7.4, 1H, C$H^{22}$).

$^{13}$C NMR (101 MHz, MeOD): 14.45 ($C^{35}H_3$), 20.92 ($C^{12}H_3$), 21.37 ($C^{11}H_3$), 23.74 ($C^{23}H_2$), 26.03 ($C^{34}H_2$), 28.44 ($C^{24}H_2$), 30.68 – 30.93 (m, C$H_2$), 32.29 ($C^{33}H_2$), 33.07 ($C^{21}H_2$), 36.33 ($C^{4}H_2$), 36.63 ($C^{5}H_2$), 39.78 ($C^{16,17}H_2$), 40.36 ($C^{9}q$), 49.76 ($C^{22}H$), 70.36 ($C^{1}H_2$), 77.35($C^{10}H$), 172.35 ($C^{18}O$), 174.18 ($C^{7}O$), 176.06 ($C^{3}O$).

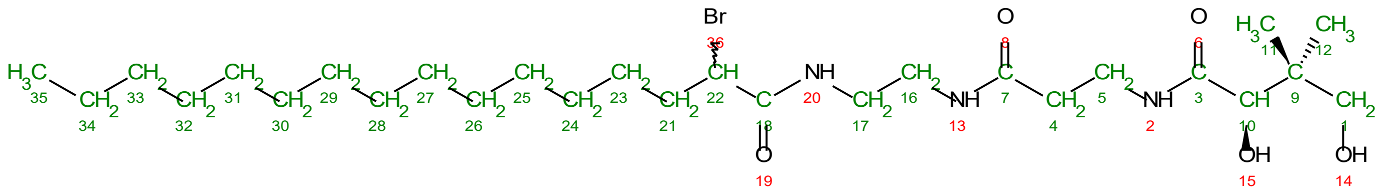

MS: $C_{27}H_{52}BrN_3O_5$ exact 577.3090, $[M+H]^+$ calculated 578.3, found 578.5 APCI (+)

### Supplementary Figures

```
Broful02171        ---------------MNKRVVITGVGMITPIGSGKEGFWSNLIAGASGVDDVSCIDTSEY 45
KustE3605          ---------------MDKRVVITGVGMITPIGSGKDVFWNALTAGESGVSEVSCIDTSGY 45
Scabro02229        ---------------MDKRIVITGLGVISPIGNQKNLFWEALISGTSGVSEVTSFDTSPF 45
Broful01969        -------------MRKRRRVVITGVGAITSLGQEKKELWSALCEGKSGIKPITSFDTTAF 47
KustD1386          -------------MHGRKRVVITGTGAITSLGREKKELWSALCNGESGIKPITSFDTTAF 47
Scabro02654        ---------------MGLRVAITGLGAITPVGYEKDLLWESLCNGKSGIANITAFDTADH 45
EcFabF             --------------MSKRRVVVTGLGMLSPVGNTVESTWKALLAGQSGISLIDHFDTSAY 46
BsFabF             --------------MTKKRVVVTGLGALSPLGNDVDTSWNNAINGVSGIGPITRVDAEEY 46
hmKAS              ---------IEGRSRLHRRVVITGIGLVTPLGVGTHLVWDRLIGGESGIVSLVGEEYKSI 51
actA               ---------------MKRRVVITGVGVRAPGGNGTRQFWELLTSGRTATRRISFFDPSPY 45
Iga11              ----------MSTATARRRVVLTGFGVISSIGTGVEEYTAGLRAGRSGARPITRFDTEGF 50
Broful02170        --------------MEKTRIVITGVSVISPIGNQKEIFWKNLANGVSGIKPITLFDVSKF 46
KustE3606          --------------MGDTRIVITGISVISPLGETKEDFWQNLINGVSGIKPVTLFDVTHF 46
Scabro02228        ---------------MNNRLVITGISVLSSIGVNKDEFWTNLTNGASGIKDITLFDASKY 45
actB               ---------MDYKDDDDKSVLITGVGVVAPNGLGLAPYWSAVLDGRHGLGPVTRFDVSRY 51
Iga12              MTTMSTATARPEATLPPGTPVITGWSAVSPYGIGRAEFAAGVRAGAKTAVKADAGLGPLP 60
                                        :** .  :. *            *

Broful02171        KVHK----GCEVK------NFKYSDYIKNGVLKKIGKGSQFAIAATRLALDDAGLDLGKT 95
KustE3605          RVHK----GCEVK------NFNYSDYIKNGVLKEIGKGSQFAIAATKLALDDAKMDLSKI 95
Scabro02229        KVHR----GCTVK------DFNYDDYTKNGSSRQIGKGSQFAVAAAKLAIEDSNIGLNNI 95
Broful01969        DVHF----GGEVS------DFEPTRWLDTKEARRLDRFAQFALIASILAVEDAELRFEGF 97
KustD1386          EVKF----GGEVS------DFNPAEWLDTKEARRLDRFSQFAIAATDIAVTDAGLNFDAI 97
Scabro02654        DVLI----AGEAA------GFDPSKWFDKKEERRLDRFSQFAVASATLALEDSGVDLDEM 95
EcFabF             ATKF----AGLVK------DFNCEDIISRKEQRKMDAFIQYGIVAGVQAMQDSGLEITEE 96
BsFabF             PAKV----AAELK------DFNVEDYMDKKEARKMDRFTQYAVVAAKMAVEDADLNITDE 96
hmKAS              PCSV----AAYVPRGSDEGQFNEQNFVSKSDIKSMSSPTIMAIGAAELAMKDSGWHPQSE 107
actA               RSQV----AAEAD------FDPVAEGFGPRELDRMDRASQFAVACAREAFAASGLDPDTL 95
Iga11              GQNT----ACEVP------DFEPGRWIHHVPLDDMGRAGQYAVAAARMAVDDAGLTEDDL 100
Broful02170        KSKQ----AGEIS------DFDAKVYLGQKGIRHIDRTSLLVSSATVLAIKDANLENNTY 96
KustE3606          NSKQ----AGEIT------DFDPKIYLGKKGIRHIDRTSLLVSSATCLAIKDANLENNTY 96
Scabro02228        NSKK----AGEIT------DFDAKEYLGKRGIRHIDRTSLLVSSAAKLAMEDANITHETY 95
actB               PATL----AGQID------DFHAPDHIPGRLLPQTDPSTRLALTAADWALQDAKADPESL 101
Iga12              SSDVCTVPG-----------FDIQEQLGPRGTAKMDRLTALALVASDGLLLDADGNRAVA 109
                           .                          .        .    .  :

Broful02171        ALERIG-VSVGTTAGEIQILEKVNYIRHKDGE-DKVDPDLFLMHPCN-NIPANVAIEFGF 152
KustE3605          ELERMG-VSVGTTAGEIQILEKVNYIRHRDGE-DKVAPDLFLMHPCN-NIPANIAIEFGL 152
Scabro02229        DPERAG-VSIGTTAGEIQILEKVNHIRHENGE-DSVDPDLFLMHPCV-NMPSNISIEFGF 152
Broful01969        DKTRGG-VIIGSGMGGLLELEAQHEILLKKGP-SRISPFLIPKLMVN-AAPAQVAIRFGL 154
KustD1386          DTARVG-VFIGSGMGGLLEFEAQHKNLLNKGP-SRVSPFMVPKLMAN-AASAHAAIRYGL 154
Scabro02654        NREKVG-AIIGTGIGGIIEIEAQHKVLLERGP-SRVSPFMITKLMAN-AAPGYIAIKFGL 152
EcFabF             NATRIG-AAIGSGIGGLGLIEENHTSLMNGGP-RKISPFFVPSTIVN-MVAGHLTIMYGL 153
BsFabF             IAPRVG-VWVGSGIGGLETLESQFEIFLTKGP-RRVSPFFVPMMIPD-MATGQISIALGA 153
hmKAS              ADQVATGVAIGMGMIPLEVVSETALNFQTKGY-NKVSPFFVPKILVN-MAAGQVSIRYKL 165
actA               DPARVG-VSLGSAVAAATSLEREYLLLSDSGRDWEVDAAWLSRHMFDYLVPSVMPAEVAW 154
Iga11              GERQAV-ITVGTTDGESHDIAVLLEQELAAGDPEAMDPVLARRINAG-RLSTVIARELRM 158
Broful02170        NGDELG-IVVGSTYGSIDSISSFDFQSLREGP-NYVNPMDFPNTVLN-APASRASIFCKA 153
KustE3606          NEDELG-IVVGSTYGSIDSISSFDFQSLREGP-NYVNPMDFPNTVIN-APASRASIFCKA 153
Scabro02228        GADELG-IVIGSTYGSIDSISSFDLEGLKEGP-TFVNPMEFPNTVLN-APASRVSIFCNA 152
actB               TDYDMG-VVTANACGGFDFTHREFRKLWSEGP--KSVSVYESFAWFYAVNTGQISIRHGM 158
Iga12              TDELTG-VVLGITMGSLENVTDFLRQSYTNARPFYVDAGRIPFGSLN-HAAGATAIRHDL 167
                             .                   .      .            .   .
```

```
Broful02171      K----GPNTIIPTACAAGNYAIGYAYDLIKFGRVDLMVAGGS-DPFSKVAYTGFARLGAI 207
KustE3605        K----GPNTIIPTACAAGNYAIGYAYDLIRFGRVDVMVAGGS-DPFSKVAFTGFARLGAI 207
Scabro02229      K----GPSTIIPTACAAGNYAIGYACDLIKLGRADIMLAGGS-DPFSKVAFVGFSRLNAI 207
Broful01969      K----GPNYALVTACTTGGNAIGEAVRTIQREDADIMVAGSSEAVVTPLTLGGFSSMKAL 210
KustD1386        R----GPNFAVVAACTTGVISIGEAVRVIQRGDADIMITGSSEAVITPLAVAGFGAMKAL 210
Scabro02654      Q----AANFSVITACASGAHAIVEAFRIVQRGEADVMFAGGTEATITPLCVSAFNNMKAL 208
EcFabF           R----GPSISIATACTSGVHNIGHAARIIAYGDADVMVAGGAEKASTPLGVGGFGAARAL 209
BsFabF           K----GVNSCTVTACATGTNSIGDAFKVIQRGDADVMVTGGTEAPLTRMSFAGFSANKAL 209
hmKAS            K----GPNHAVSTACTTGAHAVGDSFRFIAHGDADVMVAGGTDSCISPLSLAGFSRARAL 221
actA             AVGAEGPVTMVSTGCTSGLDSVGNAVRAIEEGSADVMFAGAADTPITPIVVACFDAIRAT 214
Iga11            PN---VEATTVTTACAAGNYSVGYGLDSIRSGEVDIALCGGA-DAVCRKAFALFKRFGAL 214
Broful02170      T----GLNTTISNGVTSSVDAIIYASDFLRMGRVKAVIAGGVHGLTHDIFWG-AHNSRIL 208
KustE3606        T----GLSTTISNGVTSSIDAIIYASDFLRMGRVKAVIAGGVYGLGHDIFWG-AHSSKIL 208
Scabro02228      T----GLNSTISTGTASGLDAIIYASDFLRLGRGKAVVAGGVHGLTPDVFWG-AYRSGIL 207
actB             R----GPSSALVAEQAGGLDALGHARRTIRRG-TPLVVSGGVDSALDPWGWVSQIASGRI 213
Iga12            K----GPNTTVAGGRVSGLLALNYARRLMGQGRATKYLVGSAEEFSAAHAWFEHTATASG 223
                                . .   :  .   :        . *.

Broful02171      A------PEICQPFDKNRKGMMVGEGAGMLLLESLDHAIQRN-ANIYAEIIGYGLSCDAY 260
KustE3605        A------PEICQPFDKNRKGMMVGEGAGMLVLESLESAEKRN-ANVYAEIIGYGLSCDAY 260
Scabro02229      A------PDICQPFDKNRKGLLVGEGAGMLVLESLKGALARN-ANIYAEILGYGLSCDGY 260
Broful01969      STRNDAPQKASRPFDKDRDGFVLSEGAGVVVLEEFEFARKRG-ARIYAEVLGYGLNSDAY 269
KustD1386        STHNSEPQKASRPFDKNRDGFVISEGAGIVVIEELEAAKKRG-ATIYAEILGYGMTADAY 269
Scabro02654      STRNEEPQKASRPFDRDRNGFVISEGSGIIILEEMENAKKRG-AHIYAEMLGFAMSDDGH 267
EcFabF           STRNDNPQAASRPWDKERDGFVLGDGAGMLVLEEYEHAKKRG-AKIYAELVGFGMSSDAY 268
BsFabF           STNPD-PKTASRPFDKNRDGFVMGEGAGIIVLEELEHALARG-AKIYGEIVGYGSTGDAY 267
hmKAS            STNSD-PKLACRPFHPKRDGFVMGEGAAVLVLEEYEHAVQRR-ARIYAEVLGYGLSGDAG 279
actA             TARNDDPEHASRPFDGTRDGFVLAEGAAMFVLEDYDSALARG-ARIHAEISGYATRCNAY 273
Iga11            T------PDVVRPFDKDRQGILTGEGAGILVLESLESALARG-ARIHAEVLGYGLSCDAA 267
Broful02170      SGSREGNEEISAPFDKRRNGMVIGEASALVILETLEDALKRN-APIYAEIKGYGTAFDPK 267
KustE3606        AGSNSGTLEISAPFDKRRNGMVLGEASALVILETLEDALKRN-ANIYAEVRGYGTAFDPK 267
Scabro02228      SGSRNGDIEISAPFDKRRNGFVIGEAAALLVIERLEDALERN-TKIYAEIKGYGCTFNPD 266
actB             STATD-PDRAYLPFDERAAGYVPGEGGAILVLEDSAAAEARGRHDAYGELAGCASTFDPA 272
Iga12            D-----------------PAPLLGEGCGLFLVEQAEAAERPPLAAVLSVETRVDIDDDP- 265
                                    . : .:. .:.::*    *         .         :

Broful02171      HITIPHPDGEGVVSAMKKALKSAHLQPGDVQYVSAHGTGTPANDKAETISIKKVFGNKPE 320
KustE3605        HITIPHPDGEGVISAMIKALKSANLRPEDVQYISAHGTGTPANDKAETISIKKVFGEKPE 320
Scabro02229      HITIPHPEGNGVTSAMKKALINAKIRPEDVQYISAHGTGTVANDKAETISIKKVFGEHAK 320
Broful01969      HIAAPEPHGEGAKRCMISALKDAQCNPEEVGYINAHATSTPIGDNIEMKAVKEVFGVHAP 329
KustD1386        HIAAPNSNGEGAIRCMTHALNDAGCGVETIDYINAHATSTPLGDQIEVGAIKKVFGDHAS 329
Scabro02654      HITAPHPEGAGACIAMRNALKDAKLNTEQISYINAHATSTHLGDMVEINAIKKVFGDHVN 327
EcFabF           HMTSPPENGAGAALAMANALRDAGIEASQIGYVNAHGTSTPAGDKAEAQAVKTIFGEAAS 328
BsFabF           HITAPAQDGEGGARAMQEAIKDAGIAPEEIDYINAHGTSTYYNDKYETMAIKTVFGEHAH 327
hmKAS            HITAPDPEGEGALRCMAAALKDAGVQPEEISYINAHATSTPLGDAAENKAIKHLFKDHAY 339
actA             HMTGLKADGREMAETIRVALDESRTDATDIDYINAHGSGTRQNDRHETAAYKRALGEHAR 333
Iga11            HPTAPNRD--GIARGIRLALDDAGVEQEEIDFISAHGTGTKANDKTESAAIVDVYGDAP- 324
Broful02170      MAISKDYQIDGNKRAIMSAIQDANLTLNDISYISANAYSGIYGDAMETQVMKEVFGMRAQ 327
KustE3606        MSTSKEYQTDGSIRAIRSALEDAKLTLHDISCISANAYSGVYGDAMEVRALKEVFGMRAK 327
Scabro02228      KVSRDHIDTTQGARCISIAMKDAGINAEDISYISACANSSITGDIMEARIIKDYFGDNAD 326
actB             PGSG---RPAGLERAIRLALNDAGTGPEDVDVVFADGAGVPELDAAEARAIGRVFGREG- 328
Iga12            --------GAAVTACARRALRRAGVDAGEVWAAVPCAAPTAAGRAEHEALAALVPADALS 317
                                   *:  :       :    . .          .

Broful02171      HLAISSIKSMMGHTMGAASAIEAITCALVVQNDIIPPTINYETPDPECD-LDYVPNIAR- 378
KustE3605        HLAISSIKSMIGHTMGAASVIEAIACALAVQNDIIPPTINYVTPDPECD-LDYVPNVMR- 378
Scabro02229      NLAISSIKSMLGHTMGAASAIEAIACALAIKEGVVPPTINYETKDPECD-LDFVPNVKR- 378
Broful01969      KIPISSTKSMLGHQLGASGSVEVIICALAITESVMPPTINYETPDPECMGLDFIPLVAR- 388
KustD1386        KIPVSSTKSMLGHQLGASGSVEIIICAFALNEGVIPPTINYETPDPQCSGLDFVPNEAR- 388
Scabro02654      NISVSSTKSMLGHQLGASGGVEALICSLVLDKNVMPPTINYENPDPDCDGIDFVPNEAK- 386
EcFabF           RVLVSSTKSMTGHLLGAAGAVESIYSILALRDQAVPPTINLDNPDEGCD-LDFVPHEAR- 386
BsFabF           KLAVSSTKSMTGHLLGAAGGIEAIFSILAIKEGVIPPTINIQTPDEECD-LDYVPDEAR- 385
hmKAS            ALAVSSTKGATGHLLGAAGAVEAAFTTLACYYQKLPPTLNLDCSEPEFD-LNYVPLKAQE 398
actA             RTPVSSIKSMVGHSLGAIGSLEIAACVLALEHGVVPPTANLRTSDPECD-LDYVPLEAR- 391
Iga11            -PRTVAVKSMLGHSMGAASALGAIACGLAIEHGFIPPTINHRETDPDCP-LDVVPNRAV- 381
Broful02170      LIPVSAIKSMTGECYDASGALQTIAAVMSINAHTVPPTINYKERDPDCD-LDYVANTAR- 385
KustE3606        LIPVTAIKSMTGECLDASGALQTVAAVMSLNTHTIPPTINYKEPDPECD-LGLVVERAA- 385
Scabro02228      KIPVSAIKSMTGECLDASGSLQCVAGVLAINNGIIPPTINYQEKDEECN-LDCVPNNSR- 384
actB             -VPVTVPKTTTGRLYSGGGPLDVVTALMSLREGVIAPTAGVTSVPREYG-IDLVLGEPR- 385
Iga12            RVPSMELLGDTGAASASFQIAAVLAAAEADAD---------------------------- 349
                             *    .
```

```
Broful02171      --KQKVTIALNNAHAFGGNNSCLVVKKFTGKT--- 408
KustE3605        --KQEVNIALNNAHAFGGNNSCLVIKKFTN----- 406
Scabro02229      --EMQVDIAMNNAYAFGGNNSSLILKKFTG----- 406
Broful01969      --EKKVKIALSNSFGFGGHNACIILGQV------- 414
KustD1386        --EKKIHKTLSNSFGFGGHNATIILGKV------- 414
Scabro02654      --ERSVENVMSNSFGFGGHNVSIILGKVR------ 413
EcFabF           -QVSGMEYTLCNSFGFGGTNGSLIFKKI------- 413
BsFabF           -RQE-LNYVLSNSLGFGGHNATLIFKKYQS----- 413
hmKAS            WKTEKRFIGLTNSFGFGGTNATLCIAGL------- 426
actA             --ERKLRSVLTVGSGFGGFQSAMVLRDAETAGAAA 424
Iga11            --EADVRIVQNNSSAFAGNNAVLILGTYGE----- 409
Broful02170      --VLPVKNVLINTFSRLGNNSSLIISKYKP----- 413
KustE3606        --TVPVKNVLINSSSRLGNSSSLIVSKFK------ 412
Scabro02228      --ESKVNNVLINSFSDTGNISAVIISKYS------ 411
actB             --STAPRTALVLARGRWGFNSAAVLRRFAPTP--- 415
Iga12            ---SRGRIALVCAVDRDGAVAVAVLRLIGEQR--- 378
```

**Supplementary Figure S1.** Multiple sequence alignment, showing the sequences of *amx*FabF2s and *amx*FabF$^{mut}$s as well as of canonical FabFs, ketosynthases (KS) and chain length factors (CLF). Broful02171, KustE3605, Scabro02229: *amx*FabF2s from *B. fulgida*, *K. stuttgartiensis* and *Scalindua brodae*, respectively. Broful01969, KustD1386, Scabro02654: canonical FabFs from *B. fulgida*, *K. stuttgartiensis* and *Scalindua brodae*, respectively. EcFabF: *Escherichia coli* FabF. BsFabF: *Bacillus subtilis* FabF. hmKAS: human mitochondrial ketosynthase. actA: actinorhodin ketosynthase. Iga11: *Streptomyces sp.* ketosynthase from the ishigamide biosynthesis pathway. Broful02170, KustE3606, Scabro02228: *amx*FabF$^{mut}$s from *B. fulgida*, *K. stuttgartiensis* and *Scalindua brodae*, respectively. actB: actinorhodin biosynthesis chain length factor. Iga12: *Streptomyces sp.* chain length factor from the ishigamide biosynthesis pathway. The active site cysteine and histidines of the ketosynthases are highlighted in yellow and cyan, respectively.

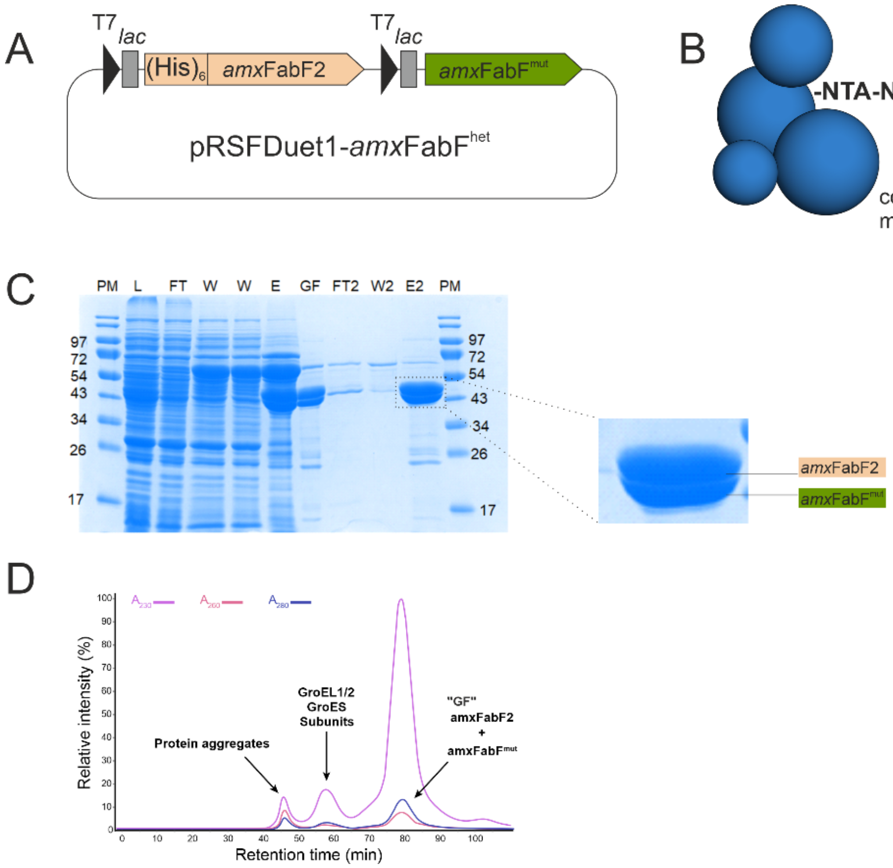


**Supplementary Figure S2. Heterodimer between *amx*FabF2 and *amx*FabF$^{mut}$.**

**A.** The two FabF homologs from Cluster 1 were expressed using a bicistronic construct, with only *amx*FabF2 displaying a His-tag. **B.** If a heterodimer is formed, then upon Ni-IMAC purification, which is based on the column material's affinity for His-tags, the non-tagged *amx*FabF$^{mut}$ should copurify with the His-tagged *amx*FabF2. **C.** Progression of a typical purification of the protein expressed using the bicistronic construct shown in **A** as shown by Coomassie-stained SDS-PAGE. PM=protein, L=loaded cell extract. FT=flow-through from 1$^{st}$ Ni-NTA column. W=wash from 1$^{st}$ Ni-NTA column. E=Elution from 1$^{st}$ Ni-NTA column. GF=result from gel-filtration (size exclusion chromatography) purification of fraction "E". This was loaded onto the 2$^{nd}$ Ni-NTA column. FT2=flow-through of the 2$^{nd}$ Ni-NTA column. W2=wash from the 2$^{nd}$ Ni-NTA column. E2=eluted fraction from the 2$^{nd}$ Ni-NTA column. The two proteins were copurified throughout the process.

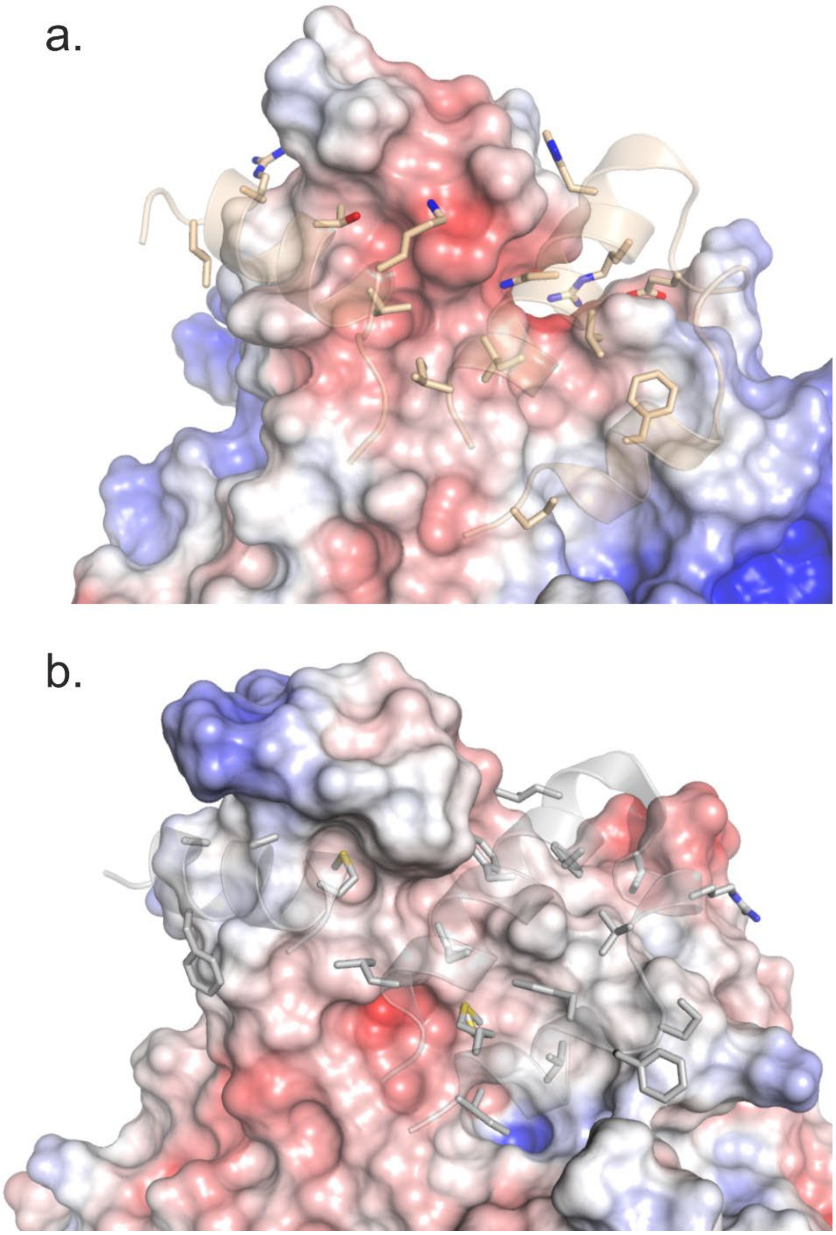


**Supplementary Figure S3. FabF dimer interfaces at the helix-turn-helix and interacting helix motifs. a.** The interface for *amx*FabF$^{het}$ is shown in panel a., with the helix-turn-helix and interacting helix motifs of *amx*FabF2 shown as cartoons and sticks, and *amx*FabF$^{mut}$ shown in surface representation, colored by electrostatic potential. On the side of *amx*FabF2, the interactions involve several positively charged side chains, whereas there is a pronounced negative charge on the interacting surface of *amx*FabF$^{mut}$. Panel **b.** shows the corresponding interface for the *E.coli* FabF homodimer, which is almost exclusively hydrophobic in nature. Electrostatic potentials are displayed on a color scale from -5 kT/e (red) to +5 kT/e (blue).

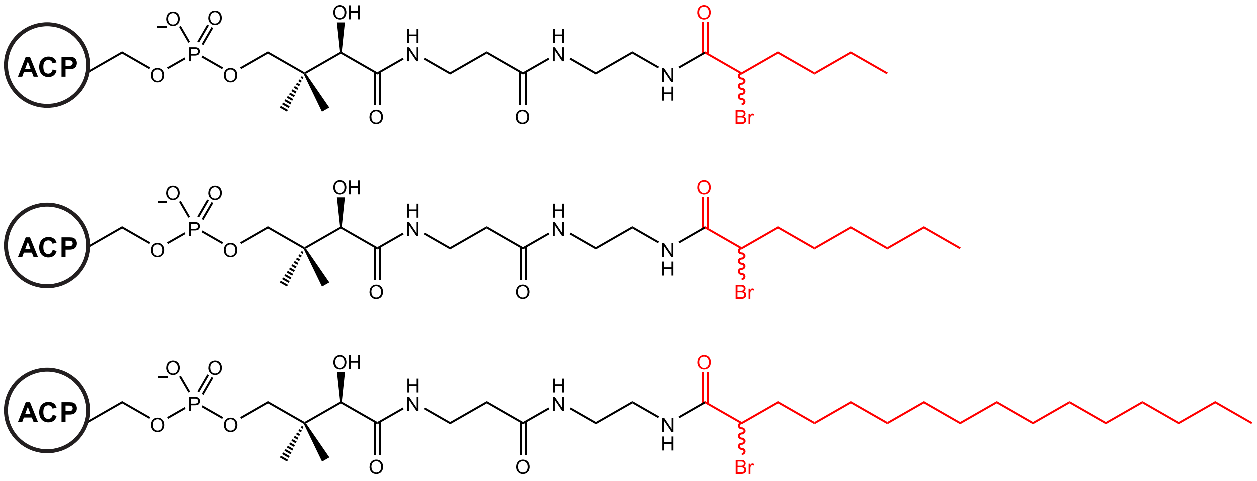


**Supplementary Figure S4. Acyl-ACP probes for crosslinking assay.** Probes were synthesized with acyl chains of 6, 8, and 16 carbons (highlighted in red) to test the chain-length specificity of the ketosynthase.

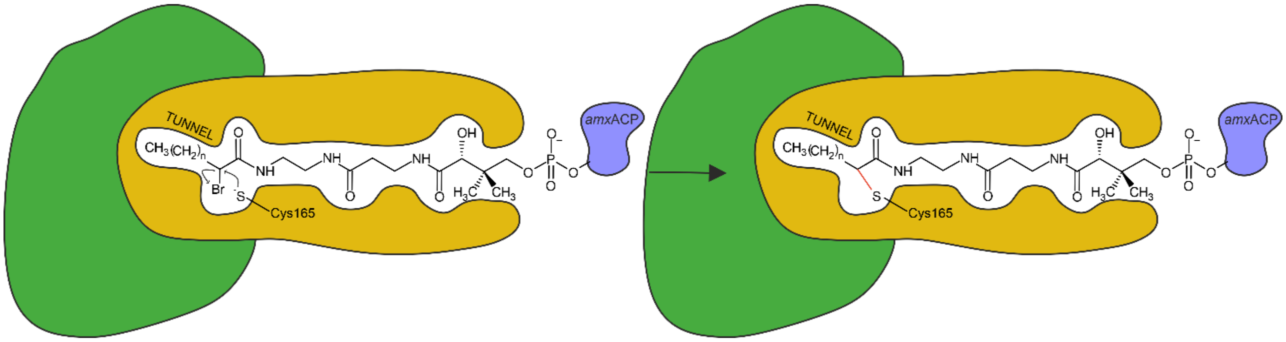


**Supplementary Figure S5. Schematic of the mechanism-based crosslinking assay.** The assay utilizes a 2-bromo-acyl-ACP probe (blue) to covalently trap the *Amx*FabF$^{het}$ complex. The probe is recognized by the heterodimer (catalytic subunit *amx*FabF2 in beige color, *amx*FabF$^{mut}$ in green), which positions the reactive 2-bromo group near the active site. The catalytic cysteine of the *amx*FabF2 subunit then performs a nucleophilic attack on the α-carbon, displacing the bromide ion and forming a stable, covalent thioether bond between the enzyme and the acyl-ACP probe.

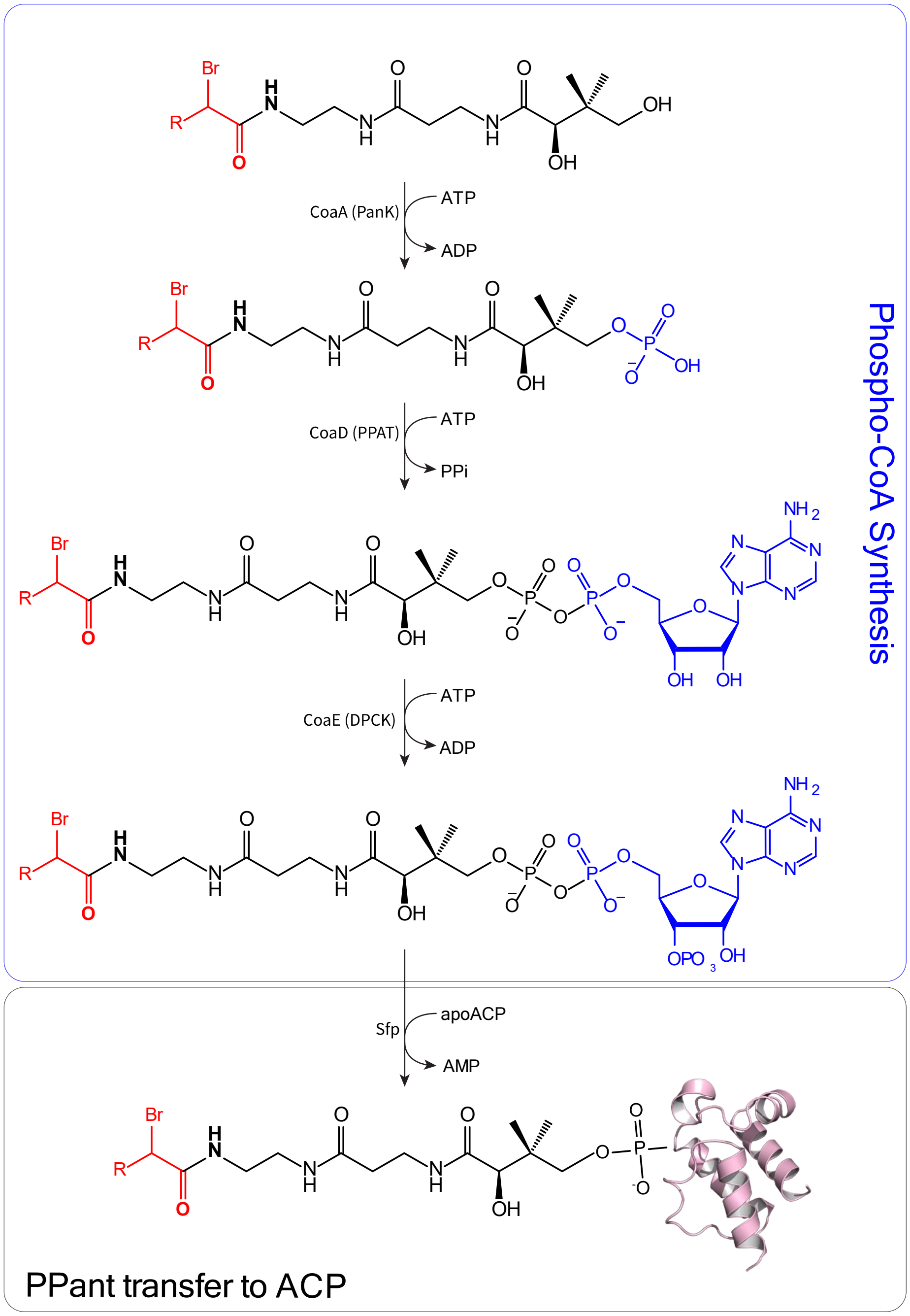


**Supplementary Figure S6. Enzymatic synthesis of the α-bromoacyl-ACP crosslinking probes.** The synthesis is a two-part, one-pot reaction. In the first stage (top panel), the chemically synthesized 2-bromo-acyl-pantetheine precursor is converted into the corresponding 2-bromo-acyl phospho-coenzyme A derivative using the *E. coli* enzyme cascade CoaA, CoaD, and CoaE. In the second stage (bottom panel), the *B. subtilis* phosphopantetheinyl transferase, Sfp, transfers the reactive 2-bromo-acyl-phosphopantetheine (PPant) arm from CoA onto a conserved serine residue of the purified apo-ACP, yielding the final, functional acyl-ACP probe.

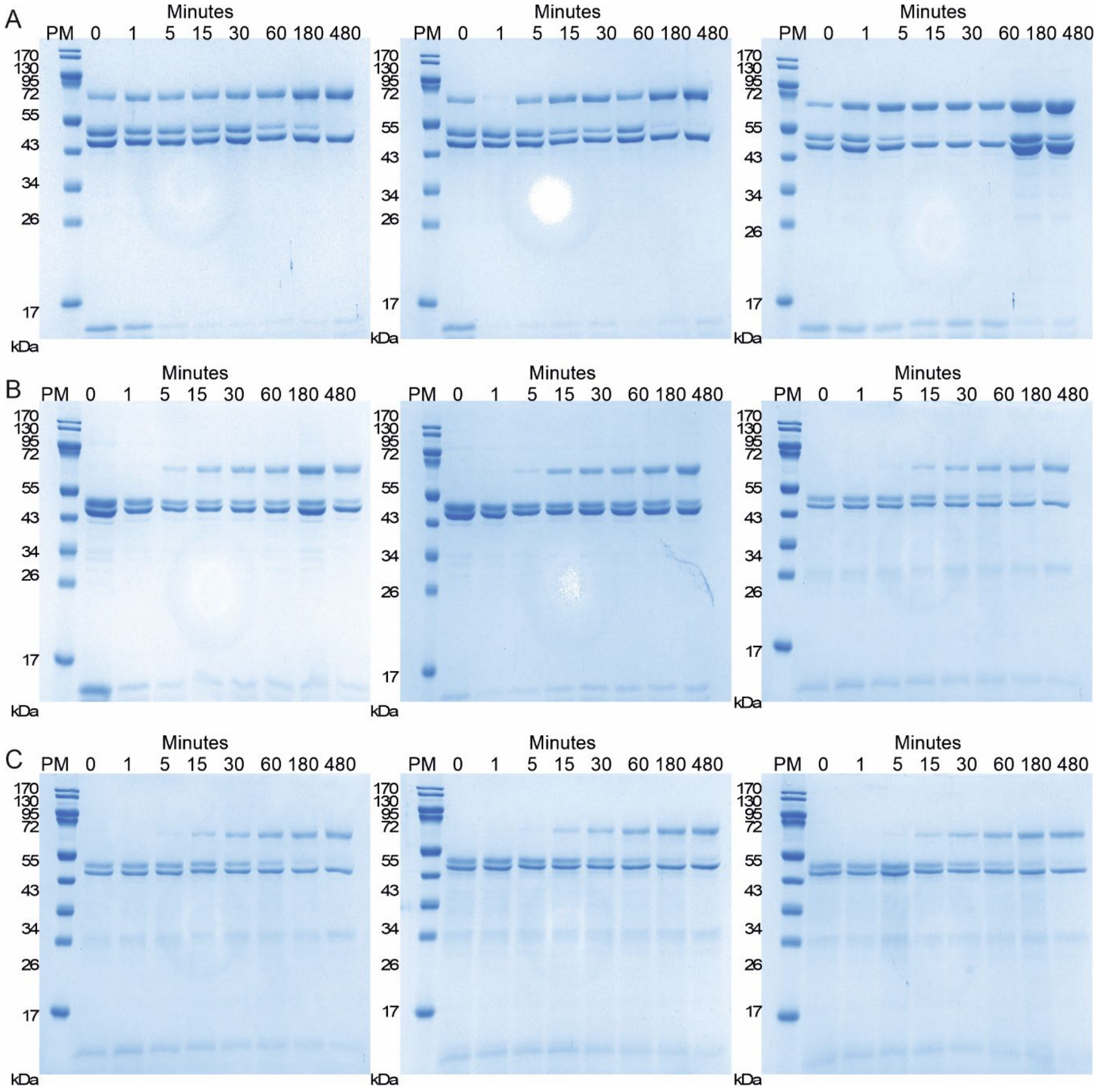


**Supplementary Figure S7. Time-Course Crosslinking Assays with $C_6$-ACP Probe**

SDS-PAGE analysis of the kinetic mechanism-based crosslinking between amxFabF$_{het}$ variants and the C6-amxACP probe (2−bromo-hexanoyl-amxACP). Reactions were sampled over 480 minutes (8 hours) at the time points indicated above the lanes (Minutes). PM denotes the protein molecular weight marker. The apparent molecular weight of the crosslinked product (amxFabF$_{het}$-amxACP adduct) is approximately 62 kDa across all panels. Unreacted protein bands, visible in all gels at approximately 45 kDa, correspond to the amxFabF2 subunit (upper band) and the amxFabF$_{mut}$ subunit (lower band). **A.** Crosslinking assay using Wild-Type, **B.** Crosslinking assay using the amxFabF$_{het}$ (V139G) mutant, **C.** Crosslinking assay using the amxFabF$_{het}$ (F394G) mutant.

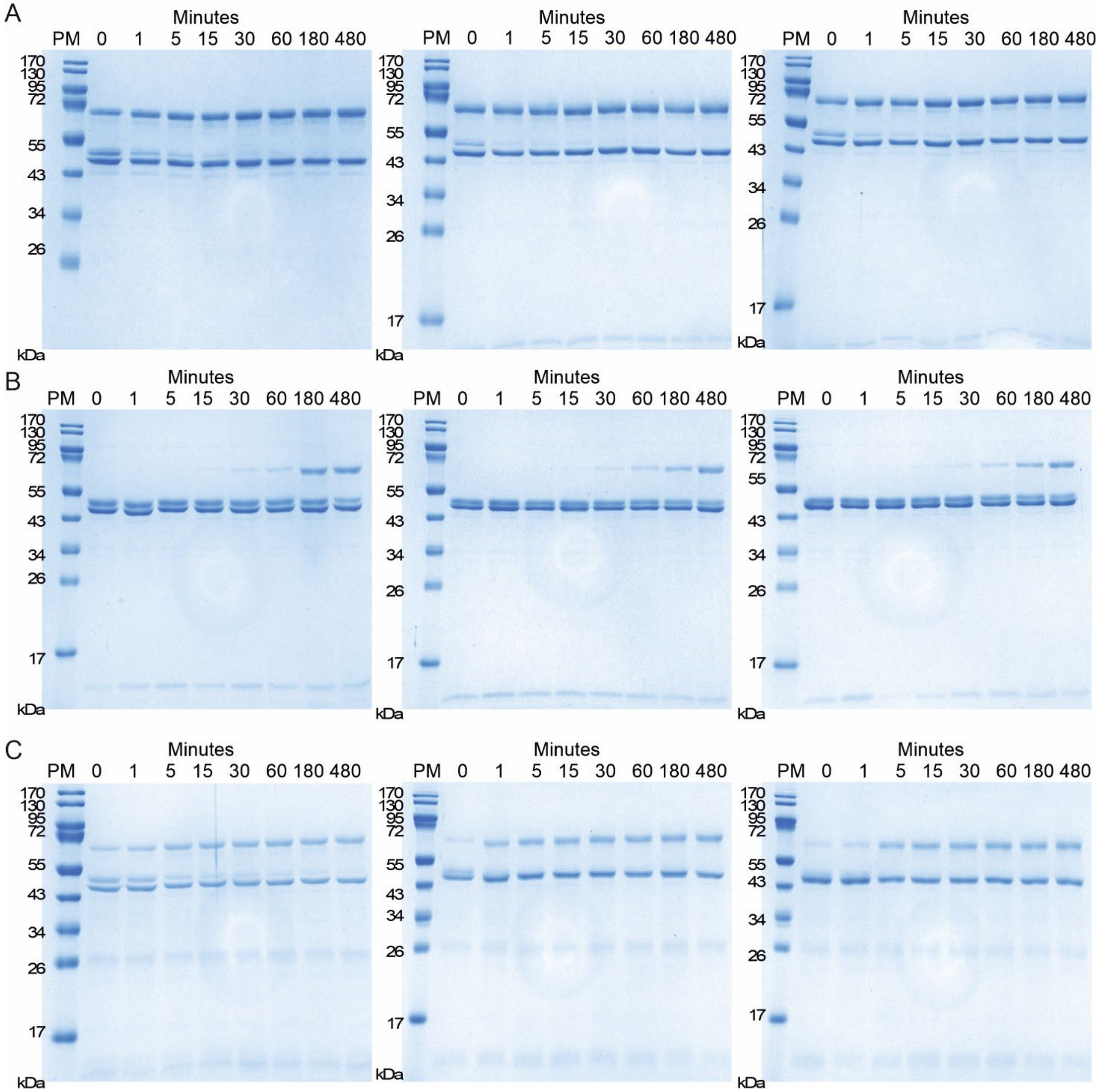


**Supplementary Figure S8. Time-Course Crosslinking Assays with $C_8$-ACP Probe**
SDS-PAGE analysis of the kinetic mechanism-based crosslinking between amxFabF$_{het}$ variants and the C8-amxACP probe (2−bromo-octanoyl-amxACP). Reactions were sampled over 480 minutes (8 hours) at the time points indicated above the lanes (Minutes). PM denotes the protein molecular weight marker. The apparent molecular weight of the crosslinked product (amxFabF$_{het}$-amxACP adduct) is approximately 62 kDa across all panels. Unreacted protein bands, visible in all gels at approximately 45 kDa, correspond to the amxFabF2 subunit (upper band) and the amxFabF$_{mut}$ subunit (lower band). **A.** Crosslinking assay using Wild-Type, **B.** Crosslinking assay using the amxFabF$_{het}$ (V139G) mutant, **C.** Crosslinking assay using the amxFabF$_{het}$ (F394G) mutant.

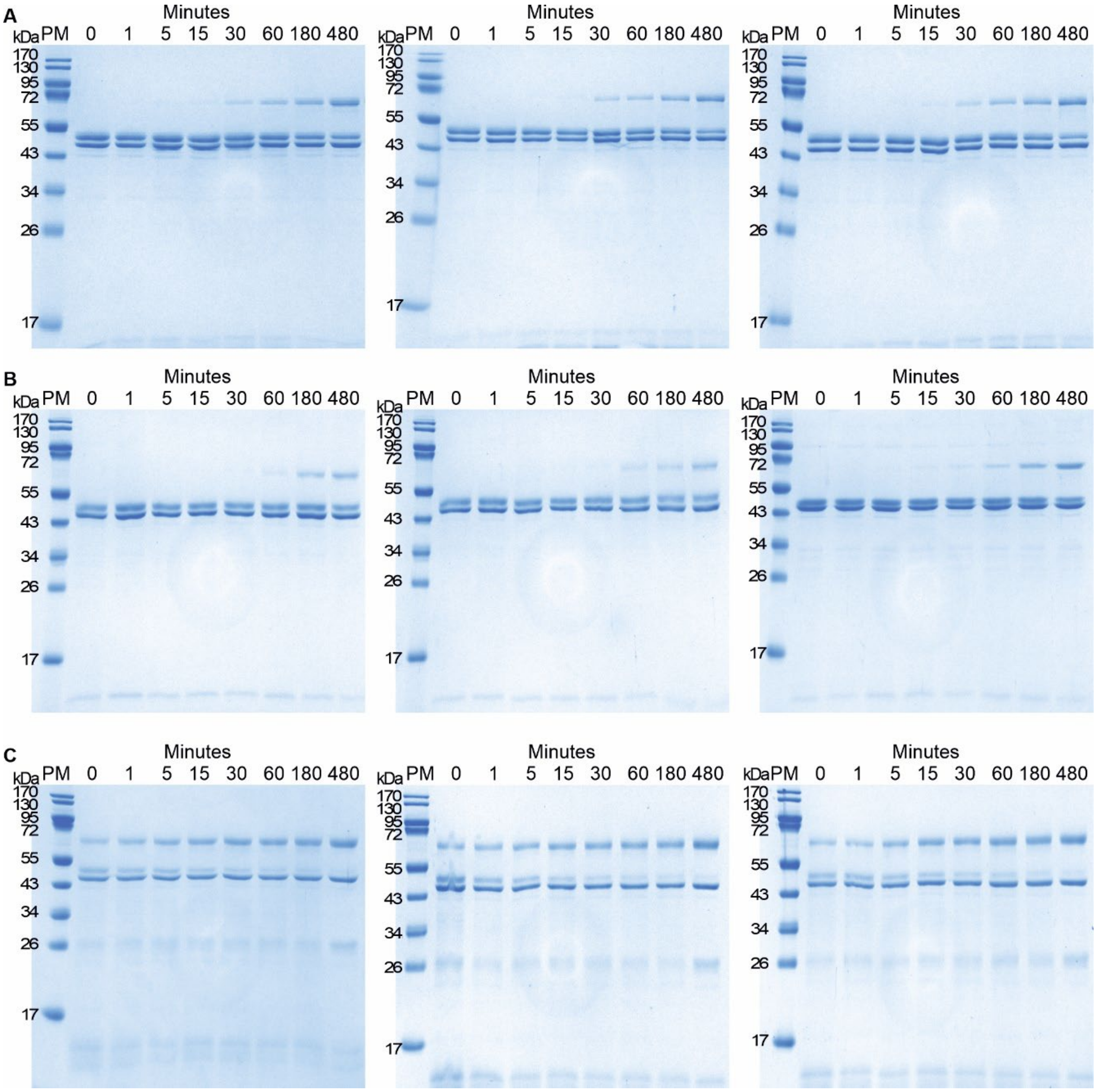


**Supplementary Figure S9. Time-Course Crosslinking Assays with C$_{16}$-ACP Probe**
SDS-PAGE analysis of the kinetic mechanism-based crosslinking between amxFabF$_{het}$ variants and the C16-amxACP probe (2−bromo-hexadecanoyl-amxACP). Reactions were sampled over 480 minutes (8 hours) at the time points indicated above the lanes (Minutes). PM denotes the protein molecular weight marker. The apparent molecular weight of the crosslinked product (amxFabF$_{het}$-amxACP adduct) is approximately 62 kDa across all panels. Unreacted protein bands, visible in all gels at approximately 45 kDa, correspond to the amxFabF2 subunit (upper band) and the amxFabF$_{mut}$ subunit (lower band). **A.** Crosslinking assay using Wild-Type, **B.** Crosslinking assay using the amxFabF$_{het}$ (V139G) mutant, **C.** Crosslinking assay using the amxFabF$_{het}$ (F394G) mutant.

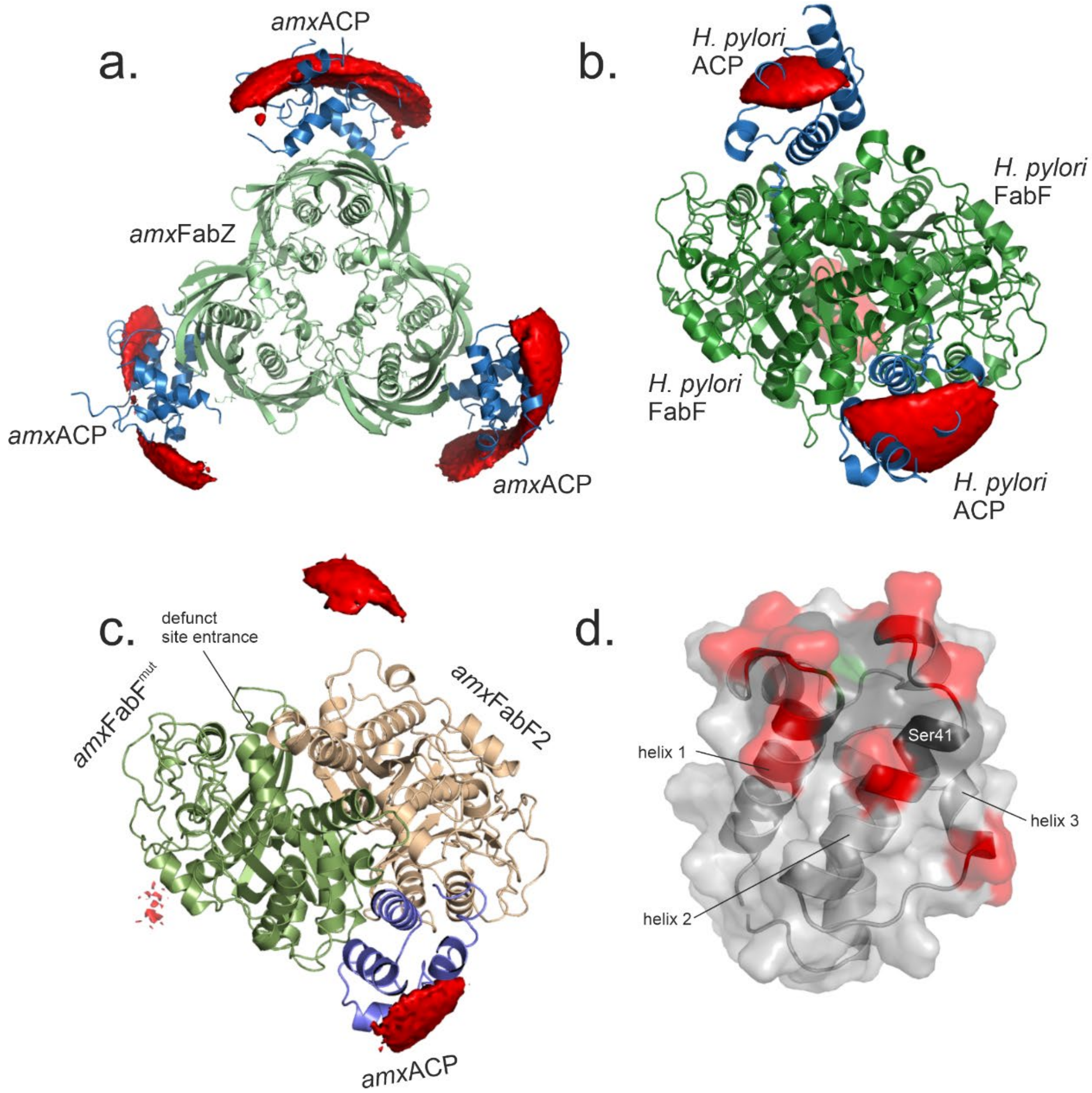


**Supplementary Figure S10. Interaction predictions with MCMAP.**

**a,b.** validation of the method for ACP/enzyme interactions; **a.** prediction of interactions between *amx*FabZ and *amx*ACP. *amx*FabZ is shown as green ribbons, the preferred locations of *amx*ACP as predicted by MCMAP are shown in red. The molecules of *amx*ACP as found in the crystal structure of the complex are shown in blue. **b.** prediction of interactions between *H. pylori* FabF and -ACP. FabF is shown as green ribbons, the preferred locations of ACP as predicted by MCMAP are shown in red. The molecules of ACP as found in the crystal structure of the complex are shown in blue. **c.** Prediction of interactions between *amx*FabF$^{het}$ and *amx*ACP. The components of the heterodimeric *amx*FabF$^{het}$ are shown in beige (*amx*FabF2) and green (*amx*FabF$^{mut}$). Preferred locations for *amx*ACP are shown in red. There is a cluster of preferred positions at the interface of the two FabF homologs in the heterodimer where the crystal structure shows that *amx*ACP is bound (blue ribbons), another on the surface of *amx*FabF2 and a minor one on the surface of *amx*FabF$^{mut}$, but none at the entrance to the defunct tunnel starting on *amx*FabF$^{mut}$. **d.** The residues on *amx*ACP predicted to be involved in interactions with *amx*FabF$^{het}$ form a negatively charged cluster on one side of the four-helical bundle. Negatively charged residues are shown in red, neutral residues in black, hydrophobic residues in green.

**Supplementary Table 1. Composition of purification buffers for *B. fulgida* and *S. brodae* constructs**

| **Buffer Step** | ***B. fulgida* Constructs (Chaperone Strain)** | ***S. brodae* (Scabro02229) (Standard Strain)** |
|---|---|---|
| **Lysis / Wash (WB)** | 100 mM Tris/HCl pH 8.0, 10 mM Imidazole, 1 mM DTE, 10% glycerol | 200 mM NaCl, 100 mM Tris/HCl pH 8.0, 1 mM Imidazole |
| **Chaperone Wash (CB)** | 50 mM HEPES pH 7.5, 300 mM KCl, 10 mM $MgCl_2$, 2 mM ATP, 10% Glycerol | **N/A** (Step omitted) |
| **Elution** | 50 mM Tris/HCl pH 8.0, 250mM Imidazole, 1 mM DTE, 10% glycerol | 200 mM NaCl, 50 mM Tris/HCl pH 8.0, 250 mM Imidazole, 1 mM DTE |
| **TEV Dialysis** | Chaperone Wash (CB) | 150 mM NaCl |
| **Gel Filtration** | 50 mM HEPES pH 7.5, 1 mM DTE | 200 mM NaCl, 50 mM HEPES pH 7.5, 1 mM DTE |

### Supplementary Table 2. Data collection and refinement statistics.

| | ***B. fulgida*** ***amx*****FabF**$^{het}$ **pdb_00009s5h** | ***B. fulgida*** ***amx*****ACP=*****amx*****FabF**$^{het}$ **pdb_00009s5i** | ***S. brodae*** ***amx*****FabF**$^{mut}$ **pdb_000030di** |
|---|---|---|---|
| **Data collection** | | | |
| Space group | $P2_12_12_1$ | $P2_12_12$ | $P2_1$ |
| *a*, *b*, *c* (Å) | 113.9, 119.9, 374.5 | 122.6, 163.2, 101.7 | 62.6, 121.4, 102.4 |
| α, β, γ (°) | 90, 90, 90 | 90, 90, 90 | 90, 102.2, 90 |
| Wavelength (Å) | 1.0000 | 0.88560 | 0.99992 |
| Resolution range (Å) | 49-2.6 (2.7-2.6) | 64-2.1 (2.15-2.1) | 50-2.2 (2.3-2.2) |
| Total no. of reflections | 729,452 (53,444) | 799,423 (50,904) | 257,824 (31,824) |
| No. of unique reflections | 157,777 (11,489) | 59,524 (3,927) | 75,298 (9,333) |
| Completeness (%) | 99.6 (99.2) | 93.9 (82.7) | 99.3 (99.1) |
| Redundancy | 4.9 (4.7) | 15.7 (15.2) | 3.4 (3.4) |
| $\langle I/\sigma(I)\rangle$ | 8.6 (1.3) | 21.1 (5.3) | 13.7 (2.6) |
| $R_{pim}$ | 0.072 (0.624) | 0.063 (0.659) | 0.066 (0.58) |
| $CC_{1/2}$ | 99.5 (48.5) | 99.9 (91.5) | 99.8 (80.6) |
| Wilson B-factor (Å$^2$) | 51.6 | 30.0 | 38.9 |
| **Refinement** | | | |
| Resolution range (Å) | 49-2.6 | 64-2.1 | 50-2.2 |
| Completeness (%) | 99.5 | 94.8 | 99.3 |
| No. of reflections | 157,676 | 117,241 | 75,290 |
| Final $R_{cryst}$ | 0.2190 | 0.1875 | 0.1748 |
| Final $R_{free}$ | 0.2666 | 0.2224 | 0.2203 |
| No. of non-H atoms | | | |
| Protein | 37,009 | 12,959 | 11,248 |
| Ligands | 143 | 104 | - |
| Water | 348 | 551 | 396 |
| R.m.s. deviations | | | |
| Bonds (Å) | 0.002 | 0.003 | 0.007 |
| Angles (°) | 0.490 | 0.620 | 0.878 |
| Average *B* factors (Å$^2$) | | | |
| Protein | 51.7 | 45.7 | 51.8 |
| Ligands | 58.0 | 55.0 | - |
| Water | 47.2 | 37.7 | 47.1 |
| Ramachandran plot | | | |
| Most favoured (%) | 97.6 | 96.8 | 97.1 |
| Allowed (%) | 2.4 | 3.2 | 2.9 |
| Outliers | 0.0 | 0.1 | 0.0 |
| Rotamer outliers (%) | 3.0 | 1.7 | 2.1 |
